# Pervasive chromothripsis drives pleural mesothelioma to impact gene function and patient survival

**DOI:** 10.64898/2026.11.04.762854

**Authors:** Kitty Sherwood, Stuart Aitken, Ryan Silk, Mihail Halachev, Stephen Fox, Devin Bendixsen, Alhafidz Hamdan, Kevin G Blyth, Colin A Semple

## Abstract

Pleural mesothelioma (PM) is a tumour type showing high structural complexity but the full spectrum of variants present and their impacts are poorly understood. We comprehensively describe the mutational landscape of mesothelioma, exploiting a large (N=443) cohort profiling the tumour genome and transcriptome in parallel. We find that the majority of candidate driver variants are mediated by copy number alterations (CNAs) and discover high frequencies of complex rearrangements across most tumours, dominated by chromothripsis (74% of tumours) and chromoplexy (59%). Chromothripsis contributes to significant CNA hotspots to delete known PM tumour suppressor genes such as *CDKN2A, NF2* and *BAP1*, thought to be among the earliest events in PM development; and the presence of chromothripsis is associated with significantly worse patient survival. We also discover multi-faceted crosstalk between the mitochondrial and nuclear genomes during PM evolution, involving alterations to both genomes, and mitochondrial expression changes associated with subtypes and chromothripsis.

## Introduction

Pleural mesothelioma (PM) is an aggressive tumour type primarily caused by asbestos exposure, with limited treatment options and poor survival rates. Global incidence is rising sharply due to continued asbestos use, at a projected rate faster than any other tumour type in men (Bizuayehu et al. 2024), us, underscoring PM as an ongoing public health challenge (Huang et al. 2023). PM has three well characterised histological subtypes. Epithelioid PM is commonest and shows tubulopapillary or solid growth, frequent BAP1 loss, and the best prognosis (Severson et al. 2020). The other subtypes are notably less common. Sarcomatoid PM is characterized by spindle cell morphology and genomic instability, while biphasic PM contains both epithelioid and sarcomatoid features and has intermediate outcome. These morphologies reflect underlying transcriptional and genomic states, with epithelioid tumours retaining mesothelial differentiation and sarcomatoid tumours showing EMT-like programs (Hmeljak et al. 2018; Severson et al. 2020). However, the development of prognostic biomarkers and more effective PM treatments is obstructed by our limited understanding of the biological processes and mutational mechanisms driving tumour development (Fennell et al, 2025).

The most recurrent single nucleotide mutations in PM affect the tumour suppressor genes (TSGs) *BAP1, CDKN2A, CDKN2B, MTAP, NF2* and *TP53*. Amplification of oncogenes is thought to be rare (Severson et al. 2020; Fennell et al. 2025), though there is evidence for recurrent amplification at specific loci such as the *TERT* gene (Mangiante et al. 2023). PM is known to be a structurally complex tumour type, but the roles of different classes of somatic variants in driving the disease remain unclear. Single nucleotide variants (SNVs), short insertions and deletions (indels), copy number alterations (CNAs) and structural variants (SVs) have all been reported as candidate drivers in studies exploiting whole exome sequencing (WES) and whole genome sequencing (WGS) (Bueno et al. 2016; Hmeljak et al. 2018; Creaney et al. 2022; Mangiante et al. 2023) but their relative importance and interactions are under studied. In addition, more recent WGS studies (Mangiante et al. 2023) have suggested that PM tumour genomes may harbour complex structural variants (CSVs), such as chromothripsis (Cortés-Ciriano et al. 2020) and extrachromosomal circular DNA (ecDNA) (Kim et al. 2020), as well as whole genome duplication (WGD) (Bielski et al. 2018). However, the full spectrum, frequencies and patterns of co-occurrence of CSVs have not been studied in PM. Beyond the nuclear genome, it is unknown whether somatic alterations also accumulate in the mitochondrial genome, though asbestos can cause mtDNA damage (Liu et al. 2013), mitochondrial metabolism has documented roles in PM (Fennell et al. 2025) and mitochondrial inhibitors have shown potential as targeted therapeutics (Gibson et al. 2026).

A critical issue in PM studies using whole exome sequencing (WES) and whole genome sequencing (WGS) has been modest cohort size, with the largest studies under-powered to interrogate the complexity and diversity of the mutational landscape. In common with many other tumour types, these studies have also used varying sequence QC, alignment and variant calling algorithms - confounding meaningful comparisons across cohorts. Thus, the driver landscape in PM remains under studied, highlighting the need for comprehensive analyses of the mutational complexity of PM and its clinical impact in larger cohorts. Here we uniformly re-process, align and analyse the raw data from four paired tumour-normal WGS (Hmeljak et al. 2018; Creaney et al. 2022; Di Genova et al. 2022; Genomics England 2025) and four tumour-normal WES (Bueno et al. 2016; Hmeljak et al. 2018; Meiller et al. 2021; Nair et al. 2023) datasets to construct the largest combined cohort to date (N=443), based upon state of the art algorithms (Watts 2024; Cortés-Ciriano et al. 2020; Deshpande et al. 2019; Imieliński and Yao 2024). These data are enhanced by matched, uniformly analysed tumour expression data for 80% of samples (N=354) to produce a comprehensive PM genomic atlas. We then describe the mutational landscape of PM to reveal a compendium of significant structural mutation hotspots, and predict candidate driver genes showing altered expression within these regions. We determine a major role for chromothripsis in driving these patterns of recurrent structural alterations, and demonstrate its association with poorer patient survival. Finally, we provide evidence for mito-nuclear crosstalk during PM evolution, suggesting links between alterations in the nuclear genome and mitochondrial metabolism.

## Results

### Complex structural variants remodel PM genome structure and function

We uniformly remapped, processed and analysed 192 PM tumour samples from four WGS studies (Supp Table 1a): The Cancer Genome Atlas (TCGA, N=57; Hmeljak et al. 2018), Creaney (N=57; Creaney et al. 2022), Di Genova (N=43; Di Genova et al. 2022), and The UK 100,000 Genomes Project from Genomics England (100kGP, N=35; Genomics England 2025). We also uniformly remapped and analysed RNA-seq for 69% (N=133) of tumours in the combined cohort. This constitutes the largest combined PM cohort with matched tumour-normal WGS to date, containing the expected proportions of the three histological subtypes (80% epithelioid, 17% biphasic, 3% sarcomatoid). The overall burdens of somatic SNVs and indels predicted to impact protein function in tumours were modest but recur in known PM driver genes (Bueno et al. 2016; Hmeljak et al. 2018), with only 8 genes (*BAP1, NF2, TP53, SETD2, LATS2, DDX3X, PBRM1* and *FBXW7;* Supp Figure 3) showing significant evidence for positive selection. Genome-wide somatic SNVs were consistent with known COSMIC single base substitution (SBS) mutational signatures (Supp Figures 7a and 7b), the majority of which are attributed to clock-like signatures associated with aging (Nik-Zainal et al. 2012; Alexandrov et al. 2013). We also identified predicted pathogenic germline variants in 21 genes from the COSMIC Cancer Gene Census (CGC) which are known to play roles in cancer (Sondka et al. 2018; Supp Table 2). The genes with the most frequent germline variants were already associated with PM risk and were often involved in the homologous recombination repair pathway, as previously reported (Belcaid et al. 2023).

We found that the mutational landscape of PM tumours is dominated by SVs and CNAs rather than by small-scale SNVs or indels. Excluding samples with low (<20%) purity to ensure high confidence variant calling, these structural changes to the genome were predicted to disrupt thousands of protein-coding genes in each sample and disproportionately involved large CNAs (median length=375Kb) (Figure 1a). As expected, *BAP1* was the most frequently altered gene and is disrupted in 84% of the cohort, often by multiple mutation types within the same tumour (Figure 1b, Supp Figure 21). The other most frequently disrupted genes mainly incurred CNA deletions and were generally not known to be associated with cancer progression, and are likely passenger events resulting from collateral damage at SV and CNA hotspots. Many of these alterations coincided with complex SVs (CSVs), which we find are remarkably common in PM, particularly chromothripsis (74% of samples), chromoplexy (59%) and templated insertion chains (TIC; 49%), impacting all four WGS subcohorts and three histological subtypes (Figure 1c). In addition, these CSVs occurred on a background of WGD (19% of samples) and homologous recombination deficiency (HRD) (8% of samples), revealing a highly complex landscape of somatic alterations. These intersecting mutational mechanisms combine to generate genomes showing high structural complexity and broad collateral damage to gene function (Figure 1a).

**Figure 1:**
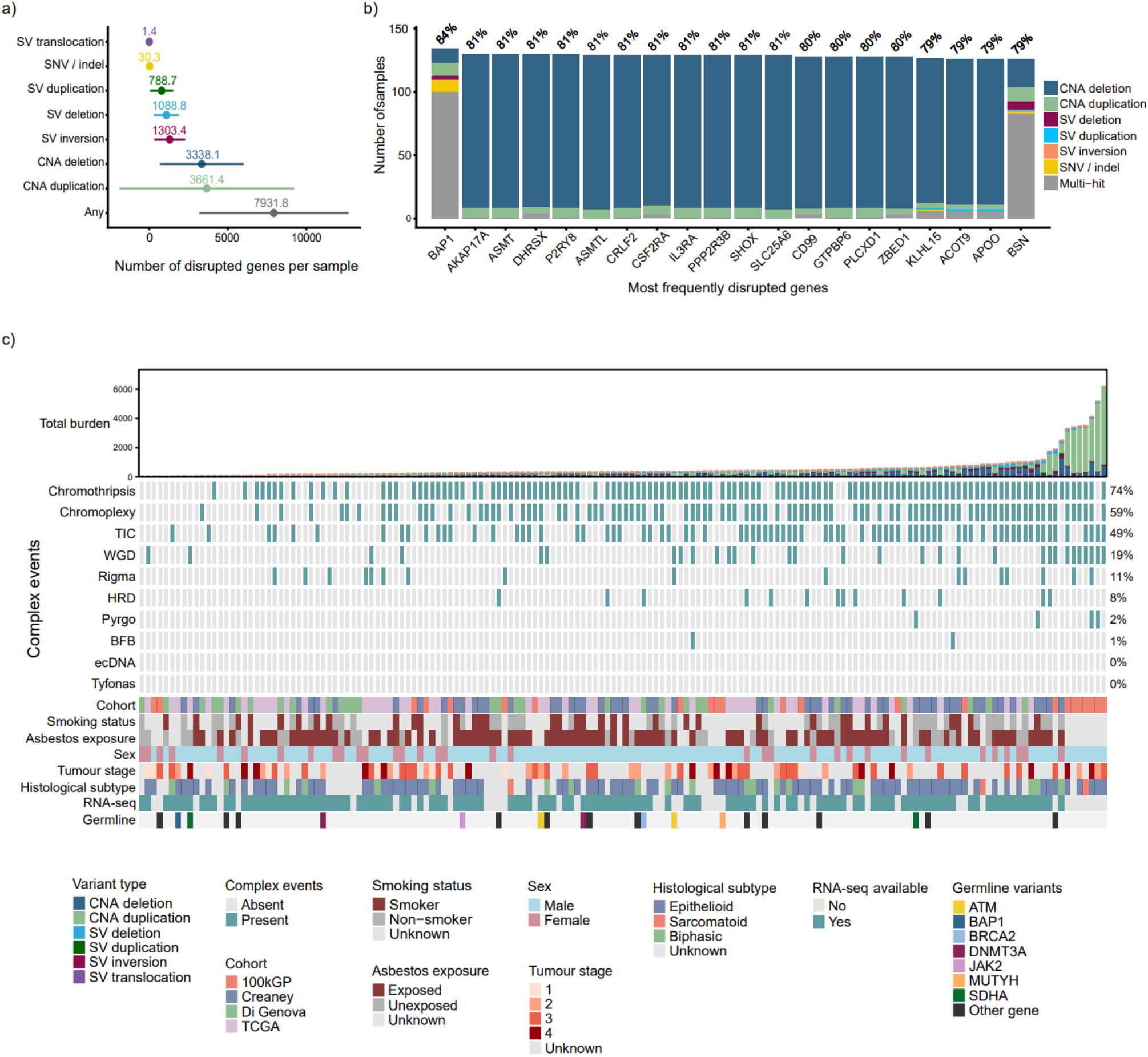
Structural complexity in the PM combined WGS cohort. a) The number of disrupted protein-coding genes predicted per sample (mean ± SD) is dominated by CNAs and SVs rather than by SNV or indel mutations. Variants predicted to disrupt genes are nonsynonymous SNVs of high/moderate functional impact and SVs or CNAs overlapping ≥1 protein-coding exon. b) The most frequently disrupted genes include the canonical PM TSG *BAP1*, and many other genes within CNA hotspots not known to be associated with cancer. c) CNAs, SVs and complex SVs dominate the mutational landscape across the combined PM WGS cohort, particularly chromothripsis, chromoplexy and templated insertions chains (TIC). These CSVs are widespread across subcohort, tumour stage, histological subtype and environmental exposures. Pathogenic germline variants are seen in genes with a known role in cancer, including known PM susceptibility genes. Genes with >1 pathogenic germline variant in the cohort are specifically labelled and genes with <=1 pathogenic variant in the cohort are displayed as “Other gene”. All data shown is for tumours with >=20% tumour purity (N=160) to ensure high quality variant calls. An unfiltered cohort overview (N=192) is provided in Supplementary Figure 4. The total burden of CNAs and SVs in the top barplot have not been filtered for predicted functional impact.

We extracted *de novo* copy number (CN) and SV mutational signatures, which correspond to known COSMIC mutational signatures (Steele et al. 2022) (Supp Figure 7c-e), including CN4, CN7, CN9 and SV9 which are associated with chromothripsis (Degasperi et al. 2020; Steele et al. 2022). We used a previously applied approach (Hart et al. 2026) to compare the mutational spectra in cancers from individuals reported to be exposed to asbestos versus unexposed in our WGS cohort (N=90 exposed versus N=31 unexposed), and found a marginal difference in *de novo* CNV signature proportional contribution in association with asbestos exposure (P=0.041, Supp Figure 8a). We also found a significant difference in SV spectra between exposed versus unexposed samples (P=0.006), which remained significant after excluding outliers (total SV burden > 2 standard deviations above the mean, P=0.014), such that the contribution of *de novo* signatures SV32C and SV32D was higher in exposed samples. Both of these signatures decompose to COSMIC SV10, which is seen in several tumour types and consists of simple SVs of a variety of sizes (Everall et al. 2026).

### Hotspots of structural alteration implicate novel candidate driver genes

The sample size of this WGS cohort allowed a comprehensive assessment of the genomic distribution of CNAs and SVs across the PM genome, which revealed a highly nonrandom distribution with abundant mutation hotspots. Of 28 significant hotspots altered by CNAs (FDR<0.05; Figure 2a, Supp Table 5), 18 were hotspots for recurrent deletions, and 10 were hotspots for recurrent duplications. The CNA hotspots were often large, with the largest a 46Mb deletion hotspot at 6q25.1, and a median size of 6Mb. The percentage of samples in the cohort with CNAs within hotspots varied widely, from 17% (Amp 6q) to 79% (Deletion 22q) (Supp Figure 9a). Deletion hotspots encompassed several regions previously reported to be recurrently deleted, including the *BAP1, NF2* and *CDKN2A* loci (Mangiante et al. 2023), among a much larger array of significant hotspots. We also identified significant CNA amplification hotspots, which are of special interest given the reported lack of amplified oncogenes in PM. In addition to CNA hotspots, we found recurrent loss or gain of whole chromosome arms across the cohort, suggesting widespread aneuploidies (Supp Table 6). Chromosome 6q was lost in 24% of the cohort and was associated with significantly improved overall patient survival (CoxPH adjusting for patient age, sex, histological subtype and stratified by subcohort; HR=0.413, P=0.005), and chromosome 14q and 14p were among several arm losses associated with worse overall survival (Supp Table 23).

**Figure 2:**
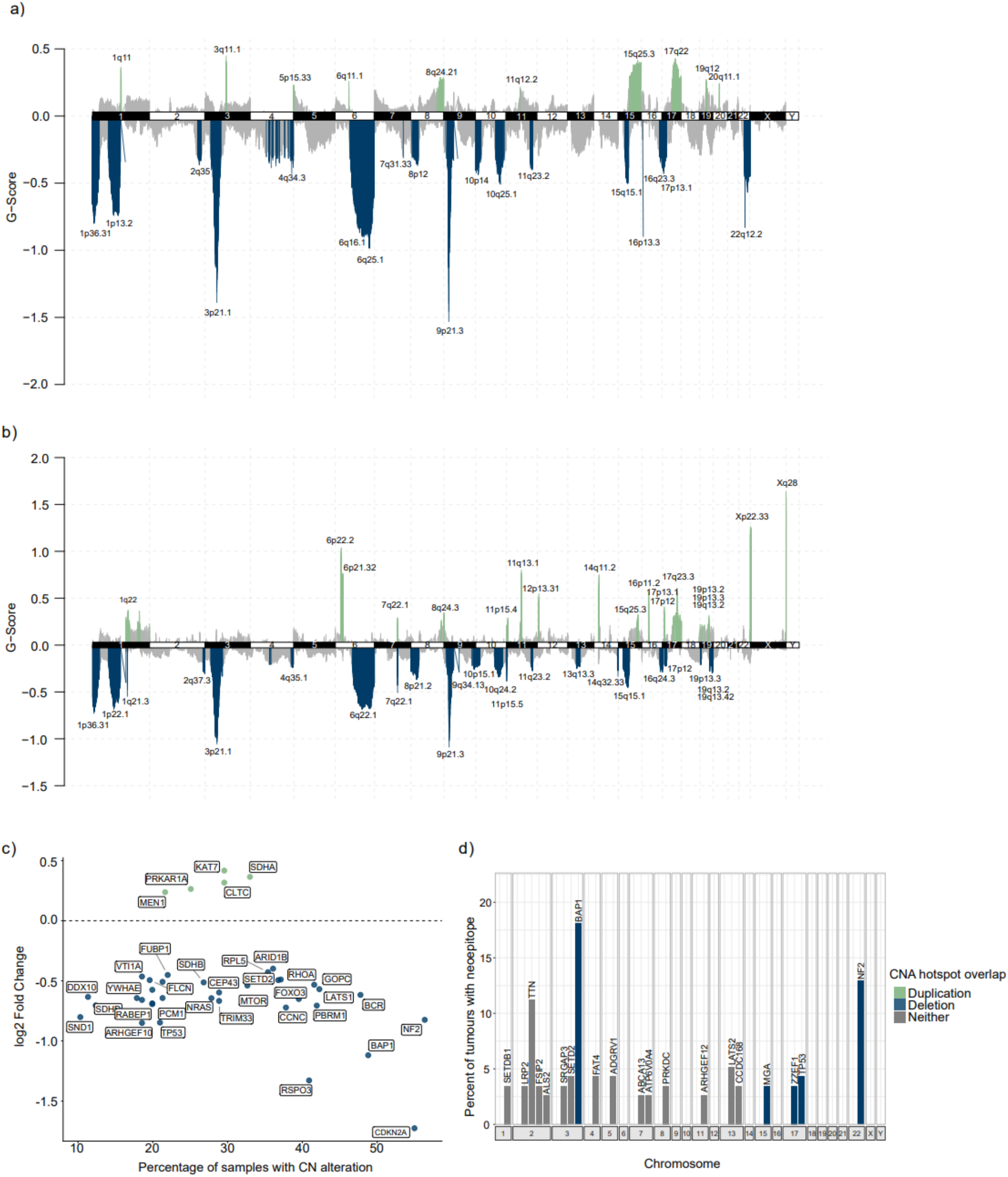
CNAs are enriched in hotspots which reveal candidate PM driver genes. (a) CNA hotspot analysis using GISTIC reveals genomic regions of significant enrichment in the WGS cohort. Duplication peaks are green and deletion peaks are in blue. G-score is a metric calculated by GISTIC considering the amplitude of the CNA and its frequency across a cohort. (b) GISTIC CNA hotspot analysis in the WES cohort shows similar patterns, and WES and WGS hotspots overlap significantly (Z=5.82, P=1×10-4, N permutations = 10,000). (c) Known CGC cancer genes within CNA hotspots showing significant differential expression across WGS and WES cohorts suggest candidate driver genes: 35 genes associated with deletion hotspots and lower expression, including several known PM TSGs, and 5 associated with duplication hotspots and higher expression. More permissive filtering of DEGs produced 105 candidate genes: 79 associated with deletions and 26 associated with duplications (Supplementary Figure 10a). (d) The top 20 genes with the highest frequency of predicted neoepitopes (filtering for the top 2% of candidates) include several SNV-mediated tumour suppressor genes within CNA deletion hotspots. Neoepitopes were predicted for 116 samples with both WGS (with >=20% purity) and RNA-seq available.

To independently validate results from the WGS cohort, we constructed a large combined WES cohort, comprising 251 uniformly analysed PM matched tumour-normal samples from five different WES studies (Supp Table 1b): Bueno (N=99; Bueno et al. 2016), TCGA (N=74; 51 of which also have WGS; Hmeljak et al. 2018), Meiller (N=18; Meiller et al. 2021) and National Cancer Institute (NCI, N=6; Nair et al. 2023). We also uniformly re-analysed the matched RNA-seq available for 177 of these samples, identically to the expression analysis for the WGS cohort. We identified 42 significant CNA hotspots in the WES cohort (Figure 2b, Supp Table 7) at FDR<0.05, comprised of 23 deletion hotspots and 19 duplication hotspots. The overall profile of CNA hotspots was similar in the WES and WGS cohorts, and permutation testing showed significant overlap (Z=5.82, P=1×10^-4^), suggesting these hotspot patterns are robust features of PM seen across independent cohorts profiled with different sequencing methods.

We identified Cancer Gene Census (CGC) genes (Sondka et al. 2018) which are known to drive cancer within many CNA deletion and duplication hotspots. Several of the CNA deletion hotspots overlapped known PM TSGs, including peaks at 3p21.1 intersecting *BAP1*, at 9p21.3 intersecting *CDKN2A,* and at 22q.12.2 intersecting NF2 (Figure 2a). We prioritised candidate CNA-mediated driver genes using a previously described approach (Ewing et al. 2025), which identifies CGC genes showing significant expression changes within hotspots, and assesses consistency across subcohorts. Based upon differentially expressed genes (DEGs with adjusted p value <=0.005; see Methods) across both WES and WGS cohorts we predicted 40 candidate CNA-mediated driver genes, of which 5 were associated with duplications and 35 with deletions (Figure 2c, Supp Tables 8 and 9). The deletion mediated list recovered most TSGs already implicated in PM, such as *BAP1, NF2, TP53, CDKN2A* and *LATS1*, supporting the validity of the approach. The 5 significant DEGs found within CNA amplification hotspots all showed significantly increased expression in response to CNA duplications (Figure 2c). These included *SDHA*, which encodes a subunit of mitochondrial complex II, within the chromosome 5p15 hotspot, and *KAT7* within the hotspot at 17q22, which is a histone acetyltransferase with oncogenic roles in colorectal cancer (H. Wang et al. 2025). More permissive filtering, based upon DEGs seen in fewer subcohorts or with significance in one of the WES or WGS combined cohorts, produced a list of 105 candidate CNA-mediated driver genes, 79 associated with deletions and 26 associated with duplications (Supp Figure 10a). Of these candidate CNA-mediated driver genes, deletions in *CDKN2A* were strongly associated with worse overall patient survival (Supp Table 23; univariate CoxPH; HR=3.900, FDR=0.001, adjusting for patient age, sex, histological subtype, chromosome 9p arm deletion and stratified by subcohort), as has been previously reported (Marshall et al. 2020). Deletions in *MLLT3,* which is often co-deleted with *CDKN2A* at chromosome 9p21.3, were also associated with worse overall survival (HR=2.593, FDR=0.027).

We identified SV hotspots for each possible SV type (Supp Figure 12, Supp Table 10) and for all combined SV breakpoints using an established approach (Imielinski et al. 2017). The majority of the 37 significant SV hotspots (median size: 9KB; range: 50Kb-8.3Mb) were found on chromosomes 3, 9 and 16, were associated with SV deletions and significantly overlapped with the coordinates of chromothripsis calls (Z=3.727, P=0.006). We then prioritised candidate driver genes within hotspots as for the CNA hotspots to reveal 6 SV-mediated candidate driver genes (*BAP1, CDKN2A, MLH1, SETD2, NCKIPSD* and *PBRM1*), all of which were also predicted as high confidence candidate CNA-mediated driver genes. Expression of all six genes was significantly downregulated in response to SV deletions, with *BAP1* and *CDKN2A* expression also significantly decreased in response to inversions (Supp Figure 12e; Supp Tables 11-13).

We also examined whether neoantigen expression might result from recurrent structural alterations. We used an established approach to predict neoepitopes in tumours with both genomic and transcriptomic data, and selected the top 2% of peptide candidates (Martínez-Jiménez et al. 2023). Tumour samples had a median of 111 candidate neoepitopes (Supp Table 14, range: 3-685) and were significantly enriched in several deletion and duplication CNA hotspots (Supp Table 15), but did not significantly overlap with chromothripsis calls (Permutation test Z=-0.515, P=0.303). *BAP1* was most frequently predicted to carry a neoepitope (present in 18% of the cohort), and several other genes frequently carrying neoepitopes were known SNV-mediated driver genes within CNA deletion hotspots (Figure 2d). However, the burden of candidate neoepitopes was not associated with histological subtype (Supp Figure 13a) or overall patient survival (CoxPH HR=0.999, P=0.414 adjusting for patient age, sex and stratified by subcohort; Supp Figure 13c).

We exploited the WGS cohort to survey the somatic mutation landscape of mtDNA in PM, which has been largely neglected in previous genomic studies, though asbestos fibres are known to damage mtDNA (Shukla et al. 2003). We found high mtDNA abundance and levels of somatic SNVs in tumour samples (Supp Figures 14 and 15; Supp Table 16), with the 13 protein coding genes encoded in mtDNA containing truncating and missense mutations in 68% of the WGS cohort samples. The highest deleterious mutation loads accumulated at particular hotspot genes (Supp Figure 14b), particularly MT-ND5 and MT-CO1, similar to the profile observed in other tumour types with high genomic instability (Ewing et al. 2025). Hotspot genes also included the mitochondrial rRNA subunits MT-RNR1 and MT-RNR2, which have been shown to accumulate deleterious SNVs under positive selection in a pan-cancer study (Boscenco et al. 2025), and we found a modest, though significant, protective effect of rRNA SNV variants on patient survival (Supp Figure 14c). The presence of mtDNA variants in rRNA subunits was also negatively associated with chromothripsis occurring in the same tumour (OR=0.3073, P=1.911×10^-3^, FDR=0.021; Supp Figure 23).

### Chromothripsis mediates structural driver alterations and is associated with poor patient survival

Complex events are extremely prevalent in PM, with ∼90% of tumours having at least one CSV called. The most common CSV type predicted was chromothripsis, with high confidence and visually validated calls present in 74% of the combined WGS cohort, and seen across all subcohorts and histological subtypes (Fig 1c). This places PM among the tumour types with the highest levels of chromothripsis seen to date, such as sarcomas and gliomas (Cortés-Ciriano et al. 2020; Espejo Valle-Inclan et al. 2025). However, in contrast to these tumour types, we found no evidence for the associated generation of ecDNAs (Fig 1c). Chromothripsis was common across all tumour stages (Supp Table 19), indicating this may be an event which occurs early in tumorigenesis. Chromothripsis events were moderately enriched in samples known to be exposed to asbestos versus unexposed, but this was not significant after multiple testing adjustment (Supp Table 19; OR=2.674, P=0.033, FDR=0.322). Chromothripsis calls varied widely in span and structure, with 50% predicted to be canonical (>=60% of segments oscillating between 2 copy number states (Cortés-Ciriano et al. 2020)) and each event involving a unique combination of CNAs and SVs. However, using PCA we could not identify associations between any of these features of chromothripsis calls with histological subtype, or clinical variables such as exposure to asbestos (Supp Figure 17), suggesting that chromothripsis is a general feature of PM driving intra-tumour heterogeneity.

Chromoplexy and templated insertion clusters (TIC) were also common (in 59% and 49% of tumours respectively), and significantly co-occur with chromothripsis within the same tumours (Figure 3b, Supp Table 19), such that 33% of tumours have very high levels of genomic instability generated through these three CSV types. Other CSV types, such as breakage fusion bridges (BFBs) and ecDNA were notably rare (Figure 1c), though they are abundant in other tumour types with high levels of genomic instability (Ewing et al. 2025; Espejo Valle-Inclan et al. 2025).

**Figure 3:**
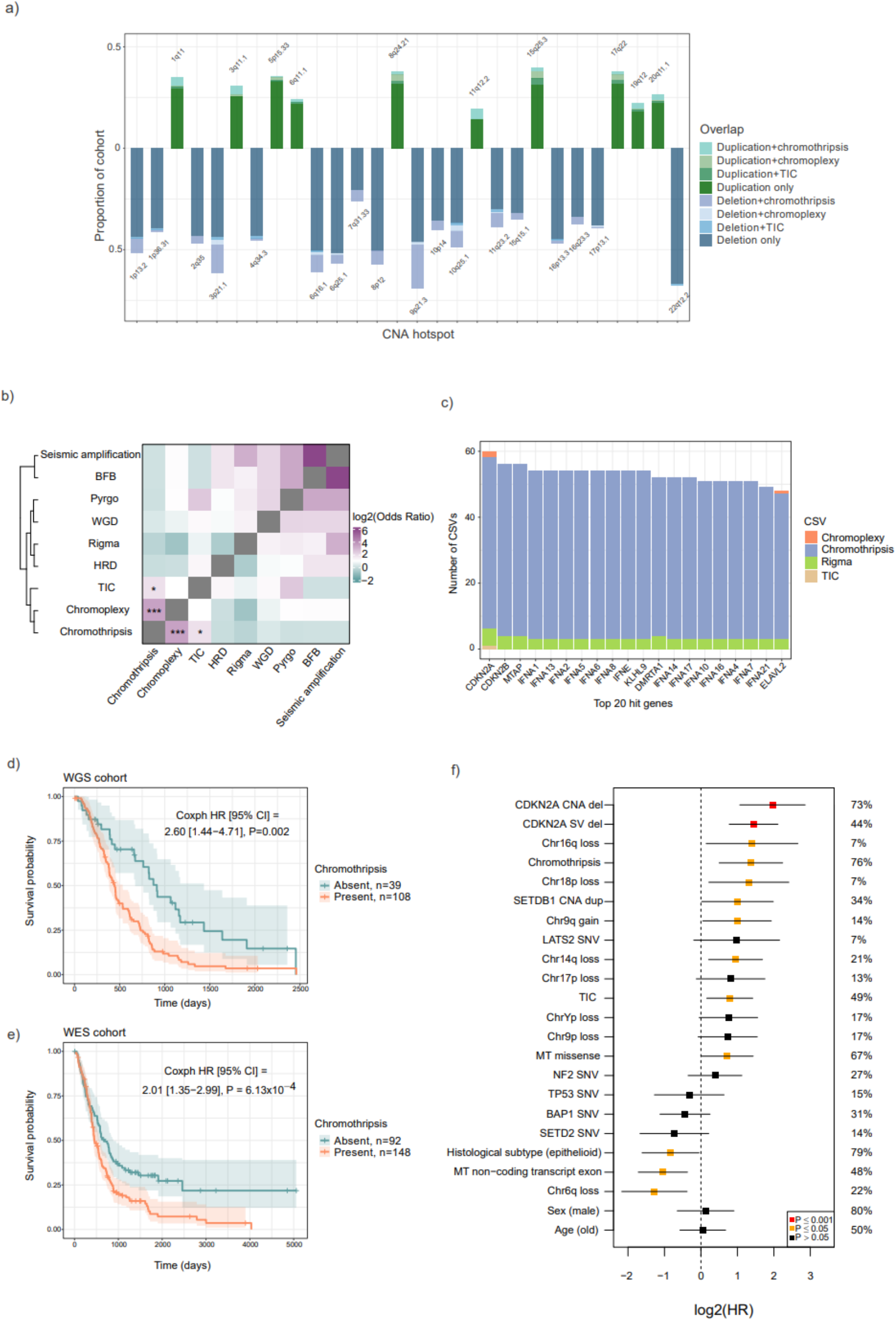
Patterns of co-occurrence between CNA hotspots and CSVs. a) CNAs within the genomic coordinates of significant hotpots with colours indicating overlap with CSV calls (chromothripsis, chromoplexy and TIC) within the same tumour sample. Chromothripsis calls significantly overlap with CNA deletion hotspots (Z=6.485, P=1×10-4, N permutations = 10,000) but not duplication hotspots (Z=0.286, P=0.407, N permutations = 10,000). b) Pairwise testing (two-tailed Fisher’s exact) and hierarchical clustering support significant co-occurrence between chromothripsis, chromoplexy and TIC; ***=FDR<0.001; *=FDR<0.05 (Supplementary Table 19). c) *CDKN2A* and the interferon alpha gene cluster are among the genes most frequently disrupted by CSVs, particularly chromothripsis. Top 50 genes hit by CSVs are provided in Supplementary Figure 18c. d) Chromothripsis in the WGS cohort is associated with worse overall survival (CoxPH, HR=2.599 [1.436-4.705], P=0.002, adjusting for patient age, sex, histological subtype and stratified by subcohort, N=105). Kaplan-Meier plot shows overall survival with 95% CI for 147 patients from WGS cohort with survival data available. e) Chromothripsis in the WES cohort based is associated with worse overall survival (CoxPH, HR=2.005 [1.347-2.985], P=6.13×10-4, adjusting for patient age, sex, histological subtype and stratified by subcohort, N=226). Kaplan-Meier plot shows overall survival with 95% CI for 240 patients from WES cohort with survival data available. f) Features retained by the 50-fold elastic net OS analysis of driver candidates, CSVs and clinical variables in the WGS cohort, with HRs from univariate CoxPH analyses, adjusting for patient age, sex, histological subtype and subcohort (N=105). Analyses of candidate CNA mediated driver genes also adjust for loss or gain of the corresponding chromosome arm. Full details provided in Supplementary Figure 32 and Supplementary Table 24. Multivariate analysis of all retained features is shown in Supplementary Figure 33.

All CSVs combined were significantly enriched on chromosomes 3, 9, 11 and 16 after adjusting for chromosome length (Supp Table 17, Supp Figure 18a), and chromothripsis calls specifically were enriched on chromosome 9 (OR=1.865, FDR=0.001, Supp Figure 18b). Remarkably, the gene most frequently disrupted by chromothripsis was *CDKN2A*, which overlapped chromothripsis calls in 38% of the WGS cohort and is a well established TSG in PM (Supp Figure 18c). *MTAP* and the interferon alpha gene cluster co-located at the 9p21.3 hotspot were also frequently disrupted by chromothripsis (Figure 3c). In comparison, chromoplexy calls were significantly enriched on chromosome 3 (OR=1.489; FDR=0.016), however chromoplexy calls intersect CGC genes at an order of magnitude less frequently than chromothripsis calls (Supp Figure 18d).

These results suggest that chromothripsis is a common mutational mechanism underlying structural driver alterations in PM. We found that chromothripsis calls significantly overlap the 18 CNA deletion hotspots in the WGS cohort (Permutation test Z=6.458, P=1×10^-4^; Supp Figure 19d), consistent with recurrent TSG loss at CNA hotspots (including at the *BAP1, CDKN2A* and *NF2* loci) mediated by chromothripsis (Figure 3a). Chromothripsis calls do not significantly overlap the 10 CNA duplication hotspots (Permutation test Z=0.286, P=0.407; Supp Figure 19e). Chromothripsis calls also significantly overlap CGC tumour suppressor genes (Z=5.398, P=1×10^-4^; Supp Figure 19b), and oncogenes (Z=2.811, P=0.002; Supp Figure 19c). Overall, it appears that chromothriptic events mediate a spectrum of driver alterations, and particularly recurrent losses of TSGs, including the canonical PM TSGs: *BAP1, CDKN2A* and *NF2*. Consistent with this, we find that the presence of chromothripsis is associated with significantly worse overall survival (CoxPH, HR=2.599 [1.436-4.705], P=0.002, FDR=0.013; adjusting for patient age, sex, histological subtype and stratified by subcohort; Figure 3d). This association remained significant after further adjusting for cancer treatment, known asbestos exposure and smoking status in the subcohort where this metadata is available (Supplementary Table 18). Increasing chromothripsis severity (number of chromosomes affected) is also associated with increasingly poorer survival (Supp Figure 20b). TIC is commonly seen alongside canonical chromothripsis (Cortés-Ciriano et al. 2020), and was also associated with worse patient survival (Supp Table 18; CoxPH;=1.739 [0.979-2.451], P=0.013, FDR=0.034), whereas chromoplexy did not have a statistically significant impact on survival (HR=1.549 [0.438-1.869], P=0.062, FDR=0.123).

WGS data is the optimal basis for prediction of chromothripsis, but previous studies (Maclachlan et al. 2021; Yu et al. 2023) have shown that this is also possible in WES profiled tumours using CNA signatures. We therefore sought to replicate the presence and clinical impact of chromothripsis in our WES cohort of 251 uniformly analysed PM tumours. We trained a binomial regression model to predict the probability of chromothripsis, using high quality calls from WGS as the outcome and copy number signatures derived from WES as predictors, based upon 51 samples from the TCGA cohort with both WGS and WES data available. Using a probability threshold of 0.7, this model achieved 79% precision, with a relatively high false positive rate, as expected given the technical limitations of WES data (Supp Table 21). We then used this model to predict chromothripsis in the entire WES cohort given a probability threshold of 0.7 (Supp Table 22), and found that 63% of the WES cohort had detectable chromothripsis, with an effect on patient survival similar to the WGS cohort (Figure 3e; CoxPH, HR=2.005 [1.347-2.985], P=6.13×10^-4^, adjusting for patient age, sex, histological subtype and stratified by subcohort).

Multi-variable regularised elastic net analysis of overall survival is effective in determining influential variables affecting survival in complex tumour genomes (Ewing et al, 2025), and we employed this strategy separately across the different classes of candidate driver events, CSVs, mtDNA mutations and aneuploidies in the WGS cohort (Supp Figure 32; Supp Table 24). Figure 3f shows the final set of 20 genomic features retained by the model (in univariate CoxPH analyses adjusting for patient age group, sex, histological subtype and stratified by subcohort). *CDKN2A* deletions were the event most strongly associated with worse patient survival. Chromothripsis was also retained in the final model and is one of the features most strongly associated with poor prognosis. Chromosome Y loss has been linked with worse progression free survival in mesothelioma (Qi et al. 2023) but without a known mechanism. We find that chromosome Yp arm loss is retained in the survival model (although just above statistical significance, HR=1.687, P=0.065), and correlates with several candidate driver events linked with poor survival, such as *CDKN2A* deletions, *TP53* SNVs and missense mtDNA mutations (Supp Figure 23). This suggests a candidate mechanism underlying the survival effects of Y loss, and possible evidence of Y loss influencing tumour evolution. In contrast, non-protein-coding mtDNA variants in rRNA or tRNA genes, loss of chromosome 6q and epithelioid histological subtype were associated with significantly improved survival. There is likely to be some redundancy among variables due to their co-occurrence. For instance, with all the retained variables combined in a single multivariate analysis (Supp Figure 33), chromothripsis remains significantly associated with worse overall survival (HR=2.981 [1.405-6.324], P=0.004), but *CDKN2A* CNA/SV deletions are no longer statistically significant (CNAs: HR=1.989 [0.856-4.622], P=0.120; SVs: HR= 2.061 [0.956-4.442], P=0.065). This suggests that chromothripsis may have some residual effect on overall survival beyond *CDKN2A* loss.

We also exploited the combined WGS/WES data to reassess the overall PM driver gene landscape, including candidate genes altered by SNVs/indels subject to positive selection (Figure 4a), as well as differentially expressed genes altered by CNA (Figure 4b) and SV hotspots (Figure 4c). These predictions suggest each tumour carries a total of 20 alterations on average that are candidate driver genes, with the majority disrupted simultaneously by large CNAs (Figure 4d). These large variants undoubtedly implicate passenger genes as well as drivers, inflating the estimated number of candidates. In addition, each tumour carries 5 further driver candidates mediated by SV hotspots, and one attributed to a SNV/indel variant. There are also distinct patterns of pairwise co-occurrence among these driver alterations, particularly for CNA/SV deletions across multiple driver genes in samples without WGD (Supp Figure 23, Supp Table 20), suggesting the possibility of genomic subtypes possessing different constellations of drivers.

**Figure 4:**
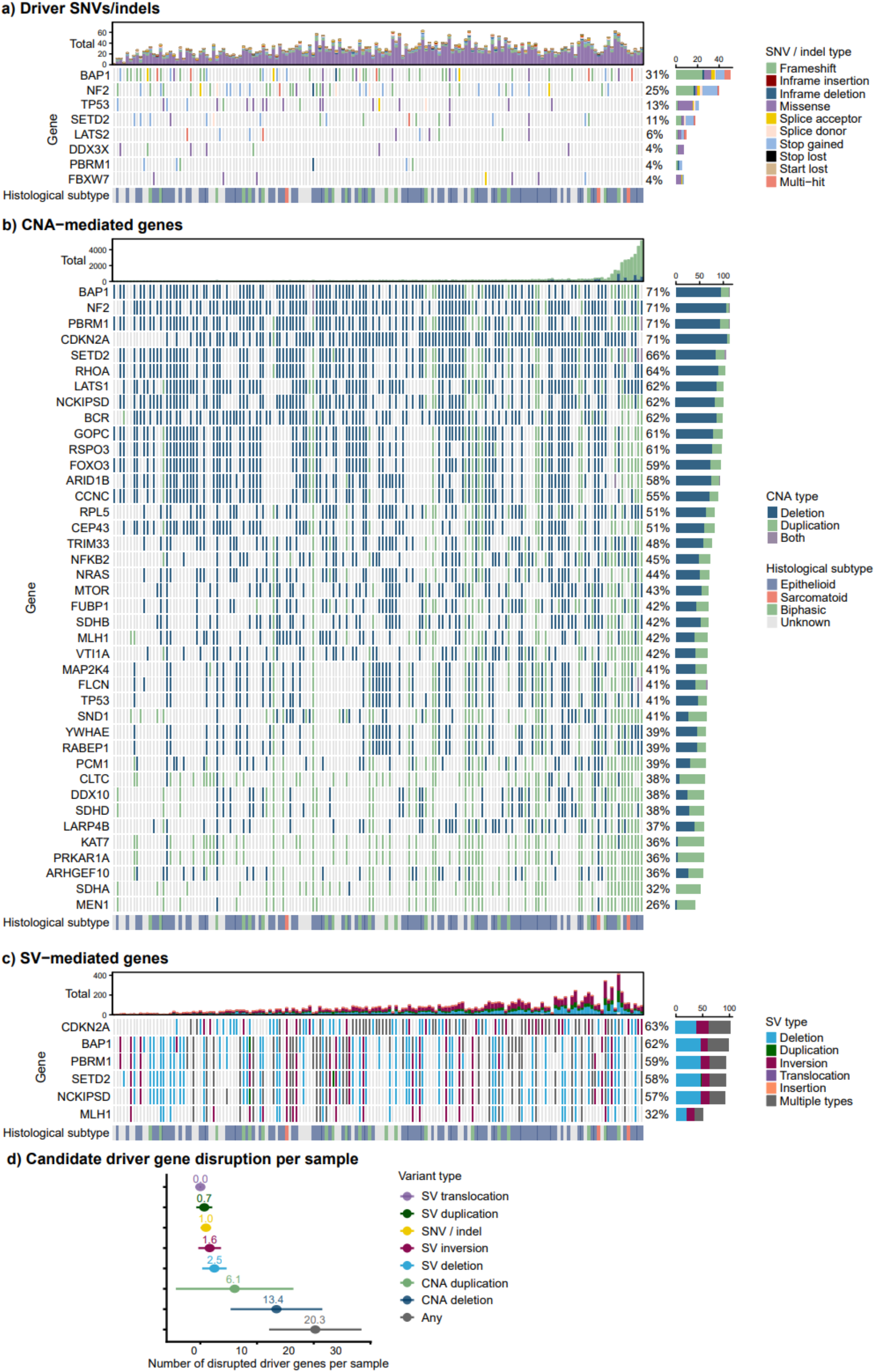
The combined driver landscape of PM. a) Candidate driver genes containing SNV/indels predicted to be under significant (q<0.05) positive selection by dNdScv. All variants shown are annotated as moderate or high functional impact from VEP. b) Candidate CNA-mediated driver genes identified as CGC genes with significant expression changes within significant CNA hotspots (see Methods). The high confidence list of 40 genes where differential expression is replicated across multiple WGS or WES subcohorts are shown. The full list of 105 candidate CNA-mediated driver genes with differential expression in a single subcohort are shown in Supplementary Figure 10b. CNAs shown are those which intersect with >=1 protein-coding exon. c) Candidate SV-mediated driver genes identified as CGC genes associated with SV hotspots with significant expression changes replicated across multiple WGS subcohorts (see Methods). SVs shown are those which intersect with >=1 protein-coding exon. d) The total number of predicted candidate driver genes disrupted per sample (mean ± SD). Variants predicted to disrupt driver genes are nonsynonymous SNVs of high/moderate impact and SVs or CNAs overlapping ≥1 protein-coding exon. In plots a-c, the samples are ordered by increasing total burden of nonsynonymous moderate/high impact SNV/indels + CNAs overlapping exons + SVs overlapping exons.

### Genomic and RNA subtypes define distinct tumour classifications associated with patient survival

In addition to histological subtype, PM subtypes have been defined by gene expression alone (de Reyniès et al. 2014; Bueno et al. 2016; Blum et al. 2019; Alcala et al. 2019) or from a combination of expression and CNAs (Hmeljak et al. 2018), while others have integrated transcriptomic, genomic, and epigenomic data (Mangiante et al. 2023). Here, we define subtypes based upon genomic and expression data as distinct layers whose interconnections we then explore.

We predicted genomic tumour subtypes based on combinatorial patterns of candidate driver events, CSVs, aneuploidies and mtDNA mutations across the WGS cohort (Methods). Using a consensus clustering approach (Lu et al. 2021) we found three genomic subtypes (CS1: n=31; CS2: n=82; CS3: n=47; Figure 5a and Supp Figure 24a-c) that were significantly associated with patient outcomes but did not correspond directly to histological subtype (2-tailed Fisher’s exact; P=0.675). Genomic subtype CS1 comprised tumour samples with WGD. Samples in CS2 were enriched for deletions at chromosome 1p36() and exclusively had loss of the chromosome 4q arm. Tumours in CS3 have fewer deletion drivers but are enriched for deletions on chromosome 6q, where 6q arm deletion is significantly protective (Figure 5e, Supp Figure 25, Supp Table 23). CS3 had significantly better overall survival compared to the other two subtypes, which remained significant after adjusting for patient age, sex, histological subtype, *CDKN2A* CNA deletion, and chromothripsis status, stratified by subcohort (Figure 5b; CoxPH CS1 vs CS3: HR=2.127, P=0.047; CS2 vs CS3: HR=2.329, P=0.014). Tumours assigned to CS3 with the best OS constitute a subset of tumours of epithelioid histological subtype.

**Figure 5:**
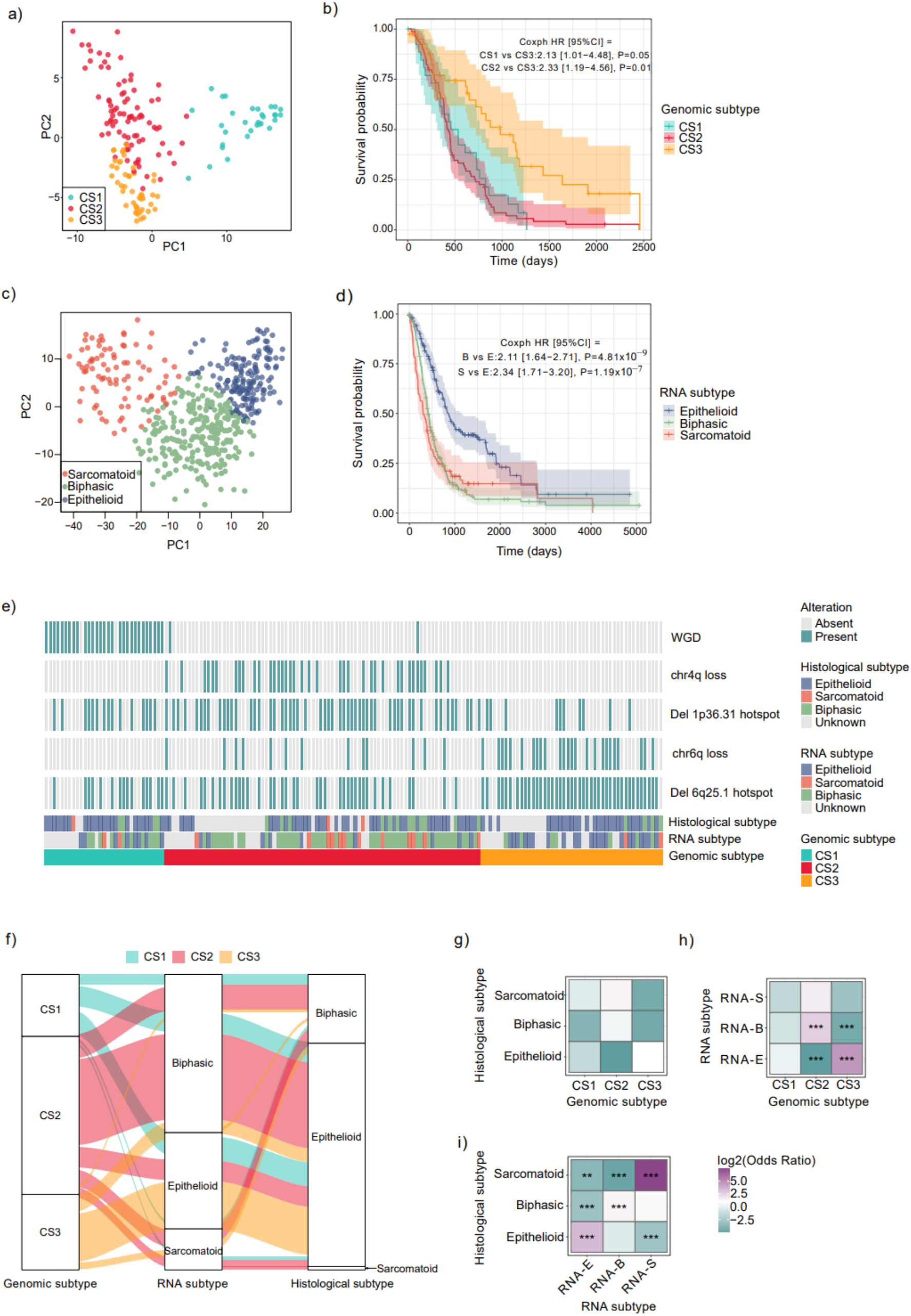
Genomic and gene expression features combine to impact overall patient survival. a) Three genomic tumour subtypes (CS1, CS2, CS3) predicted in the WGS cohort based on combinatorial patterns of CSVs and candidate driver events (Methods) occupy distinct regions in PCA. b) Genomic subtypes are significantly associated with overall patient survival. Patients with tumours in CS3 subtype have significantly improved survival compared to both CS1 and CS2 (N=160, CoxPH adjusting for patient age, sex and stratified by subcohort). c) RNA expression subtypes recovered by archetype modelling of gene expression demark distinct regions in a PCA plot of 493 samples with RNAseq from 6 WGS and WES subcohorts. d) RNA expression subtype is significantly associated with overall patient survival. Epithelioid expression subtype is associated with significantly better survival compared to both biphasic and sarcomatoid (N=493, CoxPH adjusting for patient age, sex and stratified by subcohort). e) Analysis of the most informative features of genomic subtypes (Methods, Supp Figures 24 and 25) show that CS1 is determined by WGD. CS2 is highly enriched for deletions on chromosome 1p36 and chr4q arm loss which is absent from CS3. Tumours in CS3 are also enriched for deletions on chromosome 6q. f) Genomic subtypes are strongly associated with RNA expression subtype, which in turn corresponds to histological subtype. CS3 with the best survival is enriched for tumours with epithelioid RNA expression subtype, whereas CS2 with the worst survival is enriched for tumours with biphasic RNA expression subtype. g) Genomic subtype predictions do not correspond to histological subtype (N=114, 2-tailed Fisher’s exact P=0.675). h) Genomic subtype is significantly associated with RNA expression subtype (N=116, 2-tailed Fisher’s exact P=1.56×10^-6^). i) RNA expression subtype is significantly associated with histological subtype (N=493, 2-tailed Fisher’s exact P=<2.2×10^-16^). For g-h, all statistics are from 2-tailed Fisher’s exact, with log2(Odds Ratio) on the same scale as shown. ***=FDR<0.001.

We defined three RNA subtypes using archetype modelling, as previously applied in PM (Di Genova et al. 2022; Mangiante et al. 2023), based on the 200 most variably expressed genes (Supp Table 28). These classes overlap with histological subtypes to a sufficient extent to allow us to use labels for the classes informed by histology as has been done previously (Bueno et al. 2016), namely sarcomatoid, biphasic and epithelioid RNA subtypes. As with earlier classifications, there is a strong statistical enrichment but not a one-to-one correspondence with histology (Figure 5i, Supp Figures 26a and c). The genes that the archetypes are based upon have distinct expression levels across the archetypes (Supp Figure 26b), and on mapping these subtypes to the combined RNA cohort of 493 samples, it is apparent that they are distinct by PCA (Figure 5c). This three archetype model also captures the view of continuous gradients in expression between epithelioid and sarcomatoid subtype (Blum et al. 2019), and the expected significant differences in survival between epithelioid and non-epithelioid subtypes were also seen (Figure 5d).

Genomic, RNA and histological subtypes are clearly not equivalent (Figure 5f; Supp Fig 26), and we tested the agreement between subtype classifications using Fisher’s exact and Cramer’s V (Methods). This revealed a lack of significant associations between genomic subtypes and histology (Figure 5g). In contrast, RNA subtypes show associations with both genomic subtypes and histology, supporting RNA subtypes as intermediate endotypes linking tumour genotypes and phenotypes (Figure 5h, 5i, Supp Figure 30a). Specifically, we find an enrichment of genomic subtype CS3 with the epithelioid RNA subtype and genomic subtype CS2 with biphasic RNA subtype. (The WGD-associated genomic subtype CS1 and sarcomatoid RNA subtype have no corresponding categories). More broadly, RNA subtypes show associations with genomic phenomena such as chromothripsis, chromoplexy and TIC (Figure 6d and Supp Fig 30b). Thus, genomic and RNA derived types have distinct properties yet are interrelated. This is consistent with different constellations of genomic driver events giving rise to similar expression states, while these expression states underlie histological subtypes.

**Figure 6:**
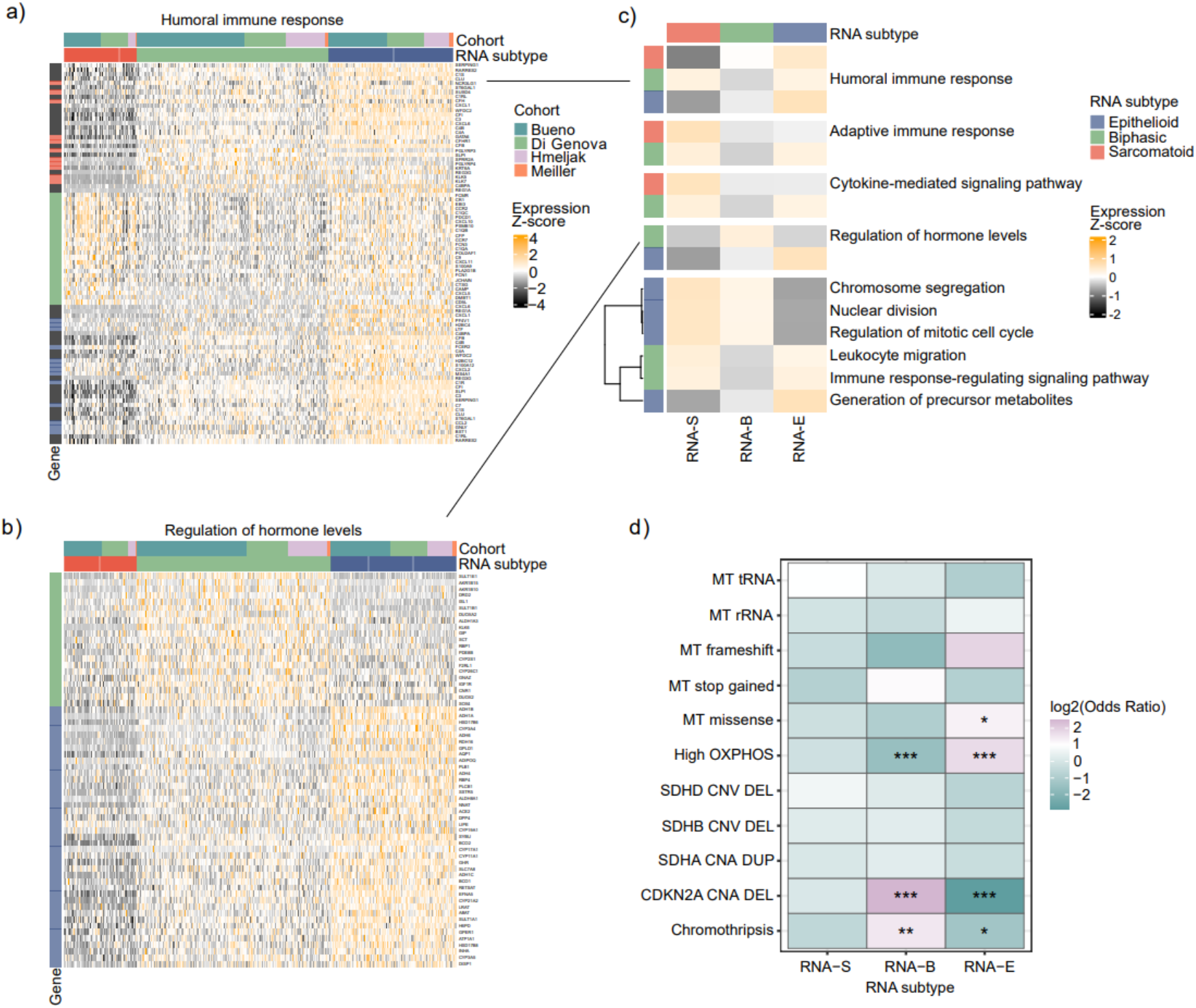
Sarcomatoid and biphasic RNA subtypes show dysregulated immunity and metabolism. Differential expression analyses show distinct patterns of gene expression between RNA subtypes in genes which are enriched for GO term functions, including a) Humoral immune response and b) Regulation of hormone levels. Each column represents a sample, with samples grouped by RNA subtype. Genes shown (rows) are those differentially expressed between RNA subtypes, with the row colour (left) indicating the RNA subtype in which it is differentially expressed. Grey rows indicate genes differentially expressed in multiple RNA subtypes. Cohort composition and expression Z-score scales are the same in both a) and b) and RNA subtypes are indicated as colours. c) Trends in differential expression between RNA subtypes (as in a) and b)) are summarized by the mean Z-score of each GO term gene set for each RNA subtype. Two sets of ‘humoral immune response’ genes have lower expression in the sarcomatoid subtype, whereas one has higher expression, while other cell cycle-related, immune and metabolomic processes show a continuous progression from epithelioid to sarcomatoid (bottom cluster). d) Pairwise tests of association (2-tailed Fisher’s exact with FDR) between RNA subtypes, chromothripsis and alterations predicted to impact mitochondrial function (N=443). High OXPHOS is defined as samples in the top quartile of gene expression for genes in the OXPHOS pathway (Supp Figure 31c).

Closer inspection of RNA subtypes reveals broad differences in pathways centred on the cell cycle, immunity and metabolism. Differential expression and pathway enrichment analyses between each RNA subtype yield sets of significantly upregulated or downregulated genes for key processes and pathways dysregulated in each subtype (Methods; Supp Table 29, Supp Figs 28 and 29). Previous studies have reported a histopathologic gradient model, where RNA and histological subtypes exist on continuous spectrum with the biphasic subtype an intermediate state on an epithelioid to sarcomatoid continuum (Bueno et al. 2016). However, we find genes in the same broad functional category may be regulated distinctly in a single RNA subtype, and that expression in the biphasic subtype is not always intermediate between the epithelioid and sarcomatoid states. For example, two sets of humoral immune response genes have lower expression in the sarcomatoid subtype, but one has higher expression (Figure 6a). Similarly, one set of hormone regulation genes has a distinctly higher expression in the biphasic subtype, whereas the second set has higher expression in epithelioid (Figure 6b). These results are consistent with a recent single cell study of PM, which defines sarcomatoid and biphasic subtypes as distinct from epithelioid, based upon the presence of a novel malignant cell state that is absent in epithelioid (Severson et al. 2026). In contrast, other dysregulated gene sets, including cell cycle-related and metabolic processes do appear to conform to the histopathologic gradient model, and show a continuous increase or decline in expression from epithelioid to sarcomatoid (Figure 6c and Supp Figure 29).

The dysregulation of mitochondrial metabolism is well established in PM, and recent work has shown that pharmacological targeting of elevated OXPHOS can suppress tumour growth *in vivo* (Gibson et al. 2026). However, it is unclear whether such strategies are equally appropriate for different subtypes. We found stark differences among RNA subtypes (Figure 6d), reflecting a spectrum of OXPHOS gene expression across samples, with highest expression in epithelioid and lowest in sarcomatoid and biphasic subtypes (Supp Figure 31c. We also examined a variety of mutation classes predicted to affect mitochondrial function across subtypes, including mtDNA mutations, SDHA duplications, *CDKN2A* deletions and chromothripsis (which often mediates *CDKN2A* deletions; Figure 3c). This revealed that the biphasic RNA subtype is associated with increased chromothripsis, *CDKN2A* deletions, and downregulated OXPHOS expression, while epithelioid tumours are relatively depleted for chromothripsis and *CDKN2A* deletions but are enriched for mitochondrial missense mutations and high OXPHOS expression (Figure 6d). This is consistent with studies showing CDKN2A loss can result in increased mitochondrial mass and impaired respiration, in addition to the well documented effects on cell cycle regulation (Tyagi et al. 2017).

## Discussion

We have shown that the genomic landscape of PM is shaped by chromothripsis, detectable in 74% of WGS tumour samples, and occurring at a higher frequency than was seen in most tumour types studied (29% of samples across 38 tumour types) using the same algorithm (Cortés-Ciriano et al. 2020). Importantly, this is an established algorithm, ShatterSeek, which combines CNA and SV calls to identify high-confidence chromothripsis events in localised regions using rigorous statistical criteria (Cortés-Ciriano et al. 2020). We also found that the CNA signatures associated with chromothripsis are detectable in the majority of available WES tumour samples, suggesting this is a pervasive feature of PM. Two previous studies observed patterns of structural variation compatible with chromothripsis in 20% (Mangiante et al. 2023) and 45% (Mansfield et al. 2019) of tumours, but both lacked statistically rigorous algorithms to confidently predict events, matched germline DNA in many cases, and the sample size to estimate overall frequency. We also found evidence for a modest, though significant, association between chromothripsis (and associated mutational signatures) and asbestos exposure, which has been hypothesised in previous literature (Gaudino et al. 2020), but requires further investigation with *in vivo* models where exposure to asbestos is precisely controlled.

The functional and clinical consequences of chromothripsis in PM are poorly studied. We have demonstrated a significant association of chromothripsis with poorer survival in independent WGS and WES cohorts, and we propose that this effect is mediated by recurrent structural alterations at known TSGs. Chromothripsis gives rise to oscillating landscapes of deletions and amplifications, and we have previously shown that these events are likely to underly a chromosome 19 amplification hotspot in ovarian cancer (Ewing et al. 2025). Remarkably, in PM, several of the CNA deletion hotspots present significantly coincided with regions subject to chromothripsis, including the *BAP1* and *CDKN2A* loci, suggesting chromothripsis drives TSG loss in many samples. In contrast, amplification hotspots did not show significant overlap with chromothriptic regions, suggesting they arise via alternative mutational mechanisms.

Previous studies have identified similar genomic regions subject to recurrent alteration but have not rigorously prioritised candidate driver genes using expression data (Bueno et al. 2016; Hmeljak et al. 2018; Mangiante et al. 2023). We have predicted 40 high confidence candidates showing significant expression changes across subcohorts in response to CNAs, including a number of previously reported PM TSGs, as well as several recurrently amplified oncogenes showing increased expression in approximately one third of patients (including *SDHA, KAT7* and *MEN1*), that have not been previously reported. *SDHA* encodes a subunit of mitochondrial complex II, and its expression is amplified in ∼20% of ovarian cancers and it is suggested this may provide survival advantage in anchorage-independent conditions (L. Wang et al. 2022), which may be relevant to the pleural fluid tumour environment. KAT7 is involved in cell-cycle regulation and the DNA damage response. Amplification has been reported in multiple cancer types and is associated with worse patient prognosis (Y. Wang et al. 2019). MEN1 expression has been shown to be upregulated in breast cancers and may be associated with dysregulation of cellular processes including proliferation (Massey et al. 2023). All three are novel candidate oncogenes in PM and may represent potential therapeutic targets. The recurrent amplification of *SDHA* might alone be expected to result in altered mitochondrial metabolism, but we also found abundant mutations in mtDNA predicted to alter mitochondrial-encoded gene functions, and evidence for mitochondrial expression changes associated with chromothripsis. This suggests a multi-faceted crosstalk between the mitochondrial and nuclear genomes during PM evolution.

Furthermore, we identified genomic PM subtypes based upon the combinations of candidate driver alterations in each tumour. Genomic subtypes showed differences in patient survival and yet did not correspond to the known PM histological subtypes, representing tumour phenotypes. However, we found that it was possible to map genomic subtypes to gene expression subtypes, which in turn relate to the histological subtypes. This is consistent with a model in which multiple different combinations of drivers can result in tumour cell lineages with similar transcriptional states, that translate into common tumour phenotypes.

Humoral immunity pathways, found to be down regulated in sarcomatoid and biphasic RNA subtypes, are driven in part by B cells, plasma cells, and in solid tumours, by mature tertiary lymphoid structures (TLS) (Leonard et al. 2026). TLS are ectopic lymphoid aggregates that can be found in chronically inflamed tissues where they can reproduce features of secondary lymphoid organs, to underpin an antitumour immune response (Leonard et al. 2026). In PM, TLS structures have been found to characterise tumours from patients with improved survival, demonstrating their potential as prognostic biomarkers (Rigutto et al. 2026). Consistent with this finding, an association between TLS presence and the epithelioid subtype was demonstrated in single cell data (Giotti et al. 2024) suggesting an underappreciated role for the B cell compartment. In addition, *MSLN* (Mesothelin) has been found to be a biomarker of PM and epithelial ovarian cancer as a result of humoral immune response (Ho et al. 2005) and has recently found to be highly expressed in epithelioid PM tumours (Severson et al. 2026). Thus, the humoral immunity genes differentially expressed between RNA subtypes reported here may have potential as prognostic biomarkers. The interplay between humoral immunity and cancer progression also suggests that modes of therapy aiming to enhance humoral immune responses might be useful in PM (Leonard et al. 2026).

Overall, these data show that PM is a tumour type which is unusually rich in chromothriptic events, generating complex rearranged genomes, and disrupting the function of hundreds or thousands of genes in each tumour. However, these chaotic patterns obscure significantly recurring hotspots of structural alterations, mediating TSG loss and apparent oncogene amplifications. Alterations in these unstable nuclear genomes are also associated with multiple alterations to mtDNA and mitochondrial expression during PM evolution, suggesting potential therapeutic avenues. Recurrent features of the genomic landscape define genomic and transcriptomic subtypes that predict patient survival differences, with implications for immunotherapeutic strategies and precision medicine.

## Methods

### WGS cohort

To maximise sample size, we have combined 157 unique PM samples from three previously published WGS cohorts (Hmeljak et al. 2018; Creaney et al. 2022; Di Genova et al. 2022). Somatic RNA-seq was also available for 133 samples. To these three studies, we have added 35 PM tumours with WGS from version 19 of the 100,000 Genomes Project (100kGP; Genomics England 2025; Supp Table 1). Any mesothelioma samples where the tumour site was not the lung pleura were excluded. All samples had matched germline with DNA extracted from whole blood. This brings the total sample size of the WGS cohort to 192 unique PM patients from the four studies, which, to our knowledge, is the largest cohort size for this tumour type with matched germline sequencing to date.

### WES validation cohort

To validate results from the WGS cohort, we created a separate WES cohort comprised of 251 participants with PM from 4 previously published studies (Bueno et al. 2016; Hmeljak et al. 2018; Meiller et al. 2021; Nair et al. 2023), selecting for tumours from pleural tissue only. All samples had matched germline with DNA extracted from whole blood. 221 tumours from this WES cohort also had RNA-seq data available. Some patients had sequencing from multiple tumours. In survival analysis and other patient-level analyses, we only included the first tumour sequenced and excluded subsequent tumour sequences.

### Sequence acquisition

The reads obtained in BAM/CRAM format were query-sorted and converted to FASTQ. Clinical annotations were also downloaded from the respective study metadata. This clinical metadata available varied between studies but included patient sex, age at diagnosis, tumour histological subtype, tumour stage, asbestos exposure and smoking status where known.

### Primary processing of WGS

We used the oncoanalyser Nextflow pipeline (v1.0.0) (Watts 2024) from the Hartwig Medical Foundation (HMF) to uniformly process and align all FASTQ files from the three previously published WGS studies. RNA-seq FASTQ files were also supplied as input to oncoanalyser in samples where available. WGS FASTQs were aligned to the HMF hg38 reference genome with alt contigs excluded using bwa-mem (v2.2.1). The 100kGP cohort bam files were re-aligned to the same HMF reference genome and then used as input to oncoanalyser.

Within oncoanalyser, germline and somatic SNVs and indels were called with SAGE (v3.3.4) and annotated with PAVE (v1.6). Germline SNVs and indels predicted as pathogenic or likely pathogenic in ClinVar (Landrum et al. 2018) were recorded (Supp Table S2). These pathogenic germline variants had no impact on patient survival (CoxPH, HR=0.891 [0.529-1.500], P=0.664). Somatic structural variants were called with GRIDSS (v2.13.2) and somatic copy number alterations (CNAs) were called with PURPLE (v4.0.2). Tumour purity was estimated using PURPLE. Samples with tumour purity below 20% were excluded from main somatic analyses, but an overview of the full cohort is provided in Supp Figure 4. Whole genome duplication (WGD) was called by PURPLE, defined as true if more than 10 autosomes had major allele copy number > 1.5. CHORD (v2.3) was also implemented within oncoanalyser to predict the probability of HR deficiency in each tumour sample, using 0.5 as the threshold score as previously described (Nguyen et al. 2020).

We found few differences between the four WGS subcohorts after uniform processing and analysis, despite separate sequencing in different geographical locations, although samples in the 100kGP cohort had the highest sequencing coverage (Supp Figure 1).

### Primary processing of WES

We applied the Nextflow pipeline Sarek (v3.3.2) to uniformly align all FASTQ files from the WES cohort to the hg38 reference genome and perform primary processing. Germline SNVs and indels were called using Strelka (v2.9.10). Somatic SNVs and indels were called using a consensus approach between Mutect2 (v2.2), Varscan (v2.4.6) and Strelka (v2.9.10), retaining variants which passed all filters and were identified by at least two variant callers. CNAs with allele-specific information were called using ASCAT (v3.2.0) (Van Loo et al. 2010) using default parameters.

Tumours in the WES cohorts also had the expected distribution of histological subtypes (74% epithelioid, 22% biphasic, 4% sarcomatoid). We find similar CGC genes with pathogenic germline variants as the WGS cohort (Supp Table 2), and the mutational landscape is also dominated by CNAs, although we are unable to call SVs with WES data (Supp Figure 6).

### Primary processing of RNA-seq

All available RNA-seq samples were processed by the Nextflow RNA Seq pipeline (v3.12.0). QC was performed by fastqc and reads trimmed by trim galore. The expression of coding and non-coding transcripts in ensembl transcriptome version GRCh38 110 was quantified by salmon (v1.10.1), and the resulting quant.sf files processed in R using tximport (Soneson et al. 2015) and DESeq2 (Love et al. 2014). Batch effects were removed by the R package ComBat_seq prior to differential expression testing. Only protein-coding genes were tested for DE, and sample purity, when available from WES or WGS data, was used as a covariate. PCA plots were used to determine batch effects and to visualise other sample level features including sex, histological subtype and RNA subtype.

### Complex SV detection in the WGS cohort

Eight categories of cSV were predicted in the WGS cohort: chromothripsis, ecDNA, breakage fusion bridges, tyfonas, pyrgo, rigma, chromoplexy and seismic amplification. ShatterSeek (Cortés-Ciriano et al. 2020) was used to call regions subject to chromothripsis (Korbel and Campbell 2013) based upon GRIDSS SV calls and PURPLE CNA calls, and following the recommended thresholds for the number of interleaved SVs, the number of adjacent segments oscillating between CN states, the number of interchromosomal translocations, the fragment joints test, the chromosomal enrichment test and the exponential distribution of breakpoints test (Cortés-Ciriano et al. 2020). Candidate ecDNAs were predicted using AmpliconArchitect (Deshpande et al. 2019) with default settings within the AmpliconSuite-pipeline (v1.3.8) to call ecDNA based upon purity and ploidy adjusted somatic CN calls from PURPLE. The resulting graph and cycle output files were processed by AmpliconClassifier (H. Kim et al. 2020) to identify amplicons with CN>4.5 and size >50kb as ecDNAs (Bailey et al. 2024). The junction-balanced genome graph (JaBbA) inference algorithm was used to generate genome graphs, based upon PURPLE calls (Hadi et al. 2020). The resulting graphs were assessed as known cSV classes within gGnome (Imieliński and Yao 2024), including breakage fusion bridges, tyfonas, pyrgo, rigma and chromoplexy. JaBba and gGnome also give predictions of ecDNA predictions which agreed with the output of AmpliconArchitect. We detected putative regions of seismic amplification using an established approach (Rosswog et al. 2021) and the default threshold for amplification (CN >= 5 for diploid samples, and CN >= 9 for samples with ploidy >2). A candidate seismic amplicon was defined as one that contains amplified CN segments linked by >=14 SV rearrangements, as recommended (Rosswog et al. 2021).

### Mutational signature fitting and extraction

COSMIC single base substitution (SBS) signatures (v3.3) were fitted to tumour samples using the sigProfilerAssignment package (v0.0.30) (Díaz-Gay et al. 2023). Mutation calls from SAGE were used as input for signature assignment in the WGS cohort and consensus mutation calls were used as input for the WES cohort.

Copy number signatures and SV signatures were extracted from tumour samples using non-negative matrix factorisation (NMF) and deconvoluted to known COSMIC copy number signatures (v3.3 for CN and v3.4 for SVs) using the sigProfilerExtractor package (v1.2.3). The maximum number of signatures was set to 15 and the number of NMF replicates was set to 100. Copy number segment data from PURPLE and SV breakpoints from GRIDSS were used as input for the WGS cohort, and ASCAT segment data as input for the WES cohort.

The counts and proportions of SBS, CNA and SV mutational signature counts were analysed in association with known exposure to asbestos using binomial GLM. We used an aggregate mutation spectrum distance permutation method (AMSD) within the R package mutspecdist (v0.3.0) (Hart et al. 2026) to compare the raw counts and proportional activity of SBS, CNA and SV mutational spectra in samples known to be exposed to asbestos versus unexposed. The number of permutations was set to 10,000. Outlier samples were defined as those with a mutation burden >2 standard deviations above the mean burden within exposed and unexposed groups.

### Chromothripsis prediction in WES cohort

Copy number signatures have been shown to accurately predict chromothripsis in multiple myeloma tumours (Maclachlan et al. 2021; Yu et al. 2023) from both WGS and WES. In the TCGA cohort, there are 51 samples with both WGS and WES data available. We therefore used the extracted *de novo* copy number signatures from the WES data of these 51 samples to create a binomial regression model to predict the probability of chromothripsis, using ShatterSeek calls from WGS as the outcome and copy number signatures derived from WES as predictors. The model performance was assessed at a range of probability thresholds (Supp Table 21). We then applied this model to the CNV signatures of the remaining 200 WES samples, using 0.7 as a probability threshold to define chromothripsis presence.

### Identification of genomic regions of recurrent copy number alteration/structural variation

Regions of the genome undergoing recurrent copy number deletion or amplification were identified using GISTIC (v2.0.23) using segmentation calls from PURPLE as input in the WGS cohort and calls from ASCAT in the WES cohort. GISTIC compares the observed frequency of alteration in the region across all samples, combined with the observed magnitude of change, to the background expectation of frequency and magnitude of copy number alteration obtained by permuting genomic regions. Sex chromosomes were included, however +1 was added to copy number for the X and Y chromosomes in males. We considered regions to be significantly recurrently deleted or amplified at a q-value < 0.05 and the region boundaries were defined to ensure 95% confidence that the reported region contained the recurrent event of interest. Broader chromosome-arm level recurrent gains and losses were also identified using GISTIC. Chromosome-arm level gains and losses were called by identifying tumours with copy number duplications or deletions spanning at least 90% of the chromosome arm. Genomic regions of significant (FDR corrected p-value < 0.05) SV enrichment in the WGS cohort were identified using the R package fishHook (v0.1) (Imielinski et al. 2017). All types of SV were considered separately in addition to overall breakpoint density. We implemented a negative binomial regression model of SV density throughout the genome split into 50kb bins with 1kb overlap, adjusting for local mutation density to ensure a controlled level of inflation. Genomic regions with low mappability (<0.9) were excluded.

### Prediction of SNV-mediated driver genes

We inferred SNV/indel driver variants across coding regions of the genome in the WGS and WES cohorts separately using dNdScv (v0.0.1.0) with default parameters to identify candidate variants based on the ratio of synonymous to non-synonymous mutations, accounting for the background mutation rate for each gene and multiple testing adjustment (qglobal cv < 0.05). The full list of driver gene candidates in WGS and WES cohorts are provided in Supp Figure 3.

### Prediction of CNV/SV-mediated driver genes

Genomic windows enriched for deletions, duplications, translocations, inversions and all breakpoints (at FDR < 0.05 by fishHook) and CNA hotspots identified by GISTIC were intersected with Cancer Gene Census (CGC) genes (v102) and with the corresponding sample level consensus SV and CNA calls respectively. The CGC (Sondka et al. 2018) is an established starting point for studies of cancer driver genes, containing 758 manually curated cancer driver genes, supported by functional validation studies and evidence of recurrent SNV/CNA mutations in tumours. CGC genes in hotspots were tested for differential expression on occurrence of the SV/CNA type driving the hotspot using DESeq2 (Love et al. 2014). Samples in which the SV/CNA intersects the CGC gene were compared with those lacking a SV/CNA of that type in the gene being tested. Differential expression was tested separately in each of the three subcohorts for which RNA-seq data was available, and in a combined, batch corrected, table of expression values combining cohorts. An adjusted p-value of 0.05 for the gene being tested in the DESeq2 results was considered significant. The log2 fold change reported is that from the combined expression analysis, with the number of cohorts with a significant change reported as a replication score.

All permutation testing to calculate statistical overlap between two sets of genomic coordinates was performed using the regioneR package (v1.30.0) (Gel et al. 2016) with 10,000 permutations.

### Prediction of neoepitopes

Candidate neoeptiopes were called and annotated using neo (v1.2.1) and we filtered for the top 2% of candidates with LikelihoodRank < 0.02 and ExpLikelihoodRank < 0.02, as previously recommended (Martínez-Jiménez et al. 2023). Neoepitope burden was calculated as the number of unique peptide sequences per tumour sample after filtering. For analyses involving genomic coordinates, neoepitope candidates were collapsed to unique variant events using SampleId, chromosome, position, reference allele and alternate allele.

### Analyses of mutual exclusivity and co-occurrence patterns across somatic alterations

Complex SVs were clustered according to their prevalence across samples using hierarchical clustering. Patterns of pairwise co-occurrence and mutual exclusivity between somatic events were using two-tailed Fisher’s exact tests.

### Calling mtDNA mutations and mtDNA copy number

Mitochondrial reads were extracted from the alignment files for each sample using Samtools (v1.12). Nuclear-encoded mitochondrial pseudogenes (NUMTs) can cause reads to mismap and generate false-positive variant calls, so we applied the RtN! algorithm to remove putative NUMT-derived reads, including those absent from NUMTs absent from the reference genome, leaving only reads attributable to genuine mtDNA (Woerner et al. 2020). The same extraction and filtering was applied to WGS and WES samples

### mtDNA variant calling in WGS samples

Variants were called on the filtered reads with VarScan2 (v2.4.4) (Koboldt et al. 2012), using the parameters --strand-filter 1, --min-avg-qual 30, --min-coverage 2, --min-reads 2, and --min-var-freq 0. Calls were then filtered to improve precision. We removed calls with strong strand bias (phred > 60), calls in known homopolymer-associated error-prone regions (positions 66-71, 300-316, 513-525, 3106-3107, 12418-12425, and 16182-16194), calls supported by fewer than 30 reads, and calls with heteroplasmy below 1% VAF.

### mtDNA variant calling in WES samples

Somatic mtDNA variants in WES samples were called following the approach of Gorelick et al. (Gorelick et al. 2021), taking the intersection of calls from two independent pipelines. In the first, GATK Mutect2 (v4.5.0.0) (McKenna et al. 2010) was run on the filtered reads from each tumour and matched-normal sample, with a minimum mapping quality of 10 and the parameters --mitochondria-mode true, --min-base-quality-score 20 and --pcr-indel-model AGGRESSIVE. Calls were filtered with GATK FilterMutectCalls (v4.5.0.0), and multi-allelic sites were split into individual records with bcftools norm (v1.21). The resulting VCFs were converted to MAF format with vcf2maf (v1.6.22). Variants were retained only if the alternative allele was supported by at least one read in each of the forward and reverse orientations. In the second pipeline, pileups were generated with a custom Python script using pysam (v0.22.1), with a minimum mapping quality of 20 and a minimum base alignment quality of 10. Variants were required to have at least two supporting reads in each orientation. Both pipelines applied the same final thresholds of ≥5% tumour VAF and ≥5 reads supporting the alternative allele, and only variants called by both were carried forward.

### mtDNA variant annotation and copy number estimation

SNVs and indels from both WGS and WES samples were annotated with Ensembl VEP (v112)with the --distance parameter set to 0, incorporating pathogenicity predictions from SIFT (Ng and Henikoff 2003) and PolyPhen2 (Adzhubei et al. 2013). Tumour and normal mtDNA copy number was estimated in WGS samples using a formula adapted from the PCAWG pan-cancer whole-genome analysis (Yuan et al. 2020).

### Defining genomic subtypes of tumours

To identify genomic subtypes of tumours, we implemented a consensus clustering approach using the R package MOVICS (v0.99.17) (Lu et al. 2021) for the whole WGS cohort (n=160, filtered for >=20% tumour purity). 10 different clustering algorithms were used: SNF, PINSPlus, NEMO, COCA, LRAcluster,IntNMF, CIMLR, MoCluster, iClusterBayes and ConsensusClustering. The ConsensusClustering distance was set to Euclidian and all other parameters were set as default.

The presence / absence of 183 genomic features were used as input (provided in Data Matrix): 6 complex SV types with cohort frequency >=5% (chromothripsis, chromoplexy, TIC, WGD, HRD and rigma), SNVs / indels in eight candidate driver genes (*BAP1, NF2, TP53, SETD2, LATS2, DDX3X, PBRM1* and *FBXW7*); 49 chromosome arms which were lost or gained in >=10% of the cohort (Supp Table 6); CNA deletions in 86 candidate driver genes (Supp Table 8); CNA duplications in 29 candidate driver genes (Supp Table 9); 8 SV candidate driver events (Supp Tables 11-13); and 7 classes of mitochondrial mutations (Supp Table 16).

We selected for k=3 to optimise the clustering accuracy and stability and to match the number of histological subtypes (Supp Figure 24a). The predicted genomic cluster subtypes were correlated with clinical information such as subcohort, tumour stage, sex and asbestos exposure. MOVICS also implements unadjusted survival analysis using log-rank tests with p-value adjustment. We further validated survival effects using CoxPH tests adjusting for covariates.

### Defining expression subtypes of tumours

RNA subtypes were defined directly in the space of gene expression using the archetypes method (Cutler and Breiman 1994; Seth and Eugster 2016).. A three archetype model based on the 200 most variably expressed genes defined classes that overlap sufficiently with histological subtypes to permit a histology-based reading of the archetypes (gene expression was transformed by variance stabilizing transformation in the DESeq2 R package). Archetype discovery used 320 samples from Bueno and Di Genova cohorts as these cohorts have the greatest number of sarcomatoid samples hence are most likely to span the PM expression space. Bueno, Di Genova, TCGA and Meiller cohorts were closely aligned in PCA space without batch correction being required and consequently the archetype model was used to predict subtypes in TCGA and Meiller cohorts. As Creaney and Nair cohorts were outlying in a sample PCA of all cohorts, a nearest neighbour method was used to annotate Creaney and Nair samples with an RNA subtype after combining and batch correcting data from all six cohorts.

In a four archetype model, we recovered the four subtypes proposed in Bueno et al. 2016,) where the biphasic class is further split into epithelioid and sarcomatoid subtypes almost perfectly (Cramer’s V statistic =0.89, p=1e-77). However, this model does not separate samples in a PCA plot and therefore to explore differential expression we considered the biphasic subtype as a single category (Figure 5c; Supp Figure 26e).

Following differential expression analysis where each subtype is compared with the other two, genes upregulated in the sarcomatoid subtype and downregulated in both biphasic and epithelioid subtypes were identified. GO term enrichment then identified processes, functions and cellular components that were enriched for sarcomatoid upregulated genes. Genes downregulated in the sarcomatoid subtype were found analogously, as were up and downregulated gene sets in the biphasic and epithelioid subtypes, yielding GO terms and KEGG pathways that have distinct dysregulation in PM subtypes (Supp Figure 28). A systematic selection was then made to identify 16 terms and pathways of interest (Supp Figure 29a)

### Analyses of mutual exclusivity, enrichment and and co-occurrence

Complex SVs were clustered according to their prevalence across samples using hierarchical clustering. Patterns of pairwise co-occurrence and mutual exclusivity between somatic events were using two-tailed Fisher’s exact tests.

When testing for associations between multinomial tumour classifications and somatic alterations, we used two-tailed Fisher’s exact tests when comparing two categories (e.g. epithelioid histology vs not epithelioid), and the Freeman-Halton test (a generalisation of Fisher’s exact) for comparing more than two categories, with Cramer’s V reporting the effect size of the association.

### Multivariable analyses of the impact of molecular factors on overall survival

The survival analyses considered 105 samples with complete overall survival time after diagnosis and histological subtype (TCGA=50; Creaney=47: DiGenova=0: 100kGP=8). The presence of: 10 classes of complex SV types (whole genome duplication, homologous recombination deficiency, chromothripsis, chromoplexy, templated insertion chains, breakage fusion bridges, ecDNA, rigma, pyrgo and seismic amplifications); SNVs / indels in eight candidate driver genes (*BAP1, NF2, TP53, SETD2, LATS2, DDX3X, PBRM1* and *FBXW7*); losses or gains of entire (>=90%) chromosome arms; CNA deletions in all 85 candidate driver genes (Supp Table 9); CNA duplications in all 26 candidate driver genes (Supp Table 10); 8 SV candidate driver events (Supp Tables 12-14); and 7 classes of mitochondrial mutations (all provided in Data Matrix) were tested for their impact on survival. Overall survival was modelled using a Cox proportional hazards regression model using the coxph method in the R package survival. The effect of each variable was examined individually and in combination using a multivariable model adjusted for age (greater than mean age at diagnosis), sex, histological subtype (epithelioid or other), and stratified by subcohort. Effect estimates with 95% confidence intervals are presented as forest plots (Figure 5g & and Supp Figures 32 & 33). Kaplan-Meier curves also show overall survival with 95% confidence intervals without adjusting for covariates (Figure 6).

To find out the combination of variables which are most important for overall survival, we then applied cross-validated (50-fold) elastic-net regularised Cox regression to perform variable selection using the cv.glmnet method in the R package glmnet. Variable selection was performed separately for each mutation category on features present in at least 5% of the cohort (Supp Fig 32), and retained features were combined in univariate analyses (Figure 3f) and then in a combined multivariate model of all retained features (Supp Figure 33).

## Data availability

WGS, WES and RNA-seq reads from previously published studies were downloaded in compressed FASTQ format from the sequencing facility or in aligned BAM/CRAM format (including unaligned reads) from the European Genome/Phenome Archive (Creaney et al. 2022 =EGAD00001007874, EGAD00001008341 and EGAD00001008447; Di Genova et al. 2022=EGAD00001007023 and EGAD00001007024; Bueno et al. 2016=EGAD00001001913 and EGAD00001001915; Meiller et al. 2021= EGAD00001007762). Data from the National Cancer Institute (NCI; Nair et al. 2023) was downloaded from dbGaP under accession phs002207.v1.p1. Data from The Cancer Genome Atlas (TCGA; Hmeljak et al. 2018) was downloaded from the Genomic Data Commons Portal (portal.gdc.cancer.gov/projects/TCGA-MESO, which in turn points to dbGaP accession phs000178).

Data from the National Genomic Research Library (NGRL) used in this research are available within the secure Genomics England Research Environment. Access to NGRL data is restricted to adhere to consent requirements and protect participant privacy. Genomic and associated clinical data from participants in the 100,000 Genomes Project was used in this research. Access to NGRL data is provided to approved researchers who are members of the Genomics England Research Network, subject to institutional access agreements and research project approval under participant-led governance. For more information on data access, visit: https://www.genomicsengland.co.uk/research

## Acknowledgements

We gratefully acknowledge the participants of the National Genomic Research Library (NGRL), whose contributions made this research possible. Secure access to the NGRL under project ID 1271 was provided by Genomics England, which delivers the NGRL in partnership with NHS England, and is wholly owned by the UK Department of Health and Social Care. The NGRL contains participants’ health data collected by the NHS as part of their care, along with samples and data from their participation in research, for which fully informed consent has been obtained. This includes genomic and clinical data provided through the NHS Genomic Medicine Service, as well as data obtained through research studies, including the 100,000 Genomes Project and the Generation Study, both of which are delivered in partnership with the NHS, and from other research cohorts involving external collaborators.

## Acknowledgements

CAS, KS, RS and SA were supported by MRC core funding to the MRC Human Genetics Unit, University of Edinburgh and MRC Programme funding MC_UU_00035/1.

## References

Adzhubei, Ivan, Daniel M. Jordan, and Shamil R. Sunyaev. 2013. “Predicting Functional Effect of Human Missense Mutations Using PolyPhen-2.” Current Protocols in Human Genetics / Editorial Board, Jonathan L. Haines … [et Al.] Chapter 7 (January): Unit7.20.

Alcala, Nicolas, Lise Mangiante, Nolwenn Le-Stang, et al. 2019. “Redefining Malignant Pleural Mesothelioma Types as a Continuum Uncovers Immune-Vascular Interactions.” EBioMedicine 48 (October): 191–202.

Alexandrov, Ludmil B., Serena Nik-Zainal, David C. Wedge, et al. 2013. “Signatures of Mutational Processes in Human Cancer.” Nature 500 (7463): 415–21.

Bailey, Chris, Oriol Pich, Kerstin Thol, et al. 2024. “Origins and Impact of Extrachromosomal DNA.” Nature 635 (8037): 193–200.

Belcaid, Laila, Birgitte Bertelsen, Karin Wadt, et al. 2023. “New Pathogenic Germline Variants Identified in Mesothelioma.” Lung Cancer (Amsterdam, Netherlands) 179 (107172): 107172.

Bielski, Craig M., Ahmet Zehir, Alexander V. Penson, et al. 2018. “Genome Doubling Shapes the Evolution and Prognosis of Advanced Cancers.” Nature Genetics 50 (8): 1189–95.

Bizuayehu, Habtamu Mellie, Abel F. Dadi, Kedir Y. Ahmed, et al. 2024. “Burden of 30 Cancers among Men: Global Statistics in 2022 and Projections for 2050 Using Population-Based Estimates.” Cancer 130 (21): 3708–23.

Blum, Yuna, Clément Meiller, Lisa Quetel, et al. 2019. “Dissecting Heterogeneity in Malignant Pleural Mesothelioma through Histo-Molecular Gradients for Clinical Applications.” Nature Communications 10 (1): 1333.

Boscenco, Sonia, Jacqueline Tait-Mulder, Minsoo Kim, et al. 2025. “Functionally Dominant Hotspot Mutations of Mitochondrial Ribosomal RNA Genes in Cancer.” Nature Genetics 57 (11): 2705–14.

Bueno, Raphael, Eric W. Stawiski, Leonard D. Goldstein, et al. 2016. “Comprehensive Genomic Analysis of Malignant Pleural Mesothelioma Identifies Recurrent Mutations, Gene Fusions and Splicing Alterations.” Nature Genetics 48 (4): 407–16.

Cortés-Ciriano, Isidro, Jake June-Koo Lee, Ruibin Xi, et al. 2020. “Comprehensive Analysis of Chromothripsis in 2,658 Human Cancers Using Whole-Genome Sequencing.” Nature Genetics 52 (3): 331–41.

Creaney, Jenette, Ann-Marie Patch, Venkateswar Addala, et al. 2022. “Comprehensive Genomic and Tumour Immune Profiling Reveals Potential Therapeutic Targets in Malignant Pleural Mesothelioma.” Genome Medicine 14 (1): 58.

Cutler, Adele, and Leo Breiman. 1994. “Archetypal Analysis.” Technometrics: A Journal of Statistics for the Physical, Chemical, and Engineering Sciences 36 (4): 338–47.

Degasperi, Andrea, Tauanne Dias Amarante, Jan Czarnecki, et al. 2020. “A Practical Framework and Online Tool for Mutational Signature Analyses Show Inter-Tissue Variation and Driver Dependencies.” Nature Cancer 1 (2): 249–63.

Deshpande, Viraj, Jens Luebeck, Nam-Phuong D. Nguyen, et al. 2019. “Exploring the Landscape of Focal Amplifications in Cancer Using AmpliconArchitect.” Nature Communications 10 (1): 392.

Di Genova, Alex, Lise Mangiante, Alexandra Sexton-Oates, et al. 2022. “A Molecular Phenotypic Map of Malignant Pleural Mesothelioma.” GigaScience 12 (giac128). 10.1093/gigascience/giac128.

Díaz-Gay, Marcos, Raviteja Vangara, Mark Barnes, et al. 2023. “Assigning Mutational Signatures to Individual Samples and Individual Somatic Mutations with SigProfilerAssignment.” Bioinformatics (Oxford, England) 39 (12): btad756.

Espejo Valle-Inclan, Jose, Solange De Noon, Katherine Trevers, et al. 2025. “Ongoing Chromothripsis Underpins Osteosarcoma Genome Complexity and Clonal Evolution.” Cell 188 (2): 352–370.e22.

Everall, Andrew, Avraam Tapinos, Aliah Hawari, et al. 2026. “Comprehensive Repertoire of the Chromosomal Alteration and Mutational Signatures across 16 Cancer Types.” Nature Genetics 58 (3): 570–81.

Ewing, Ailith, Alison Meynert, Ryan Silk, et al. 2025. “Divergent Trajectories to Structural Diversity Impact Patient Survival in High Grade Serous Ovarian Cancer.” Nature Communications 16 (1): 5586.

Fennell, Dean A., Yoshitaka Sekido, Paul Baas, et al. 2025. “Pleural Mesothelioma.” Nature Reviews. Disease Primers 11 (1): 56.

Gaudino, Giovanni, Jiaming Xue, and Haining Yang. 2020. “How Asbestos and Other Fibers Cause Mesothelioma.” Translational Lung Cancer Research 9 (Suppl 1): S39–46.

Gel, Bernat, Anna Díez-Villanueva, Eduard Serra, Marcus Buschbeck, Miguel A. Peinado, and Roberto Malinverni. 2016. “regioneR: An R/Bioconductor Package for the Association Analysis of Genomic Regions Based on Permutation Tests.” Bioinformatics (Oxford, England) 32 (2): 289–91.

“Genomics England.” 2025. In The National Genomic Research Library. Preprint, October 18. https://figshare.com/articles/dataset/GenomicEnglandProtocol_pdf/4530893/8.

Gibson, Victoria, Joanna Dzialo, Terri Messier, et al. 2026. “Preclinical Characterization and Phase 1 Clinical Testing of Targeting Mitochondrial Peroxiredoxin 3 in Cancer.” Nature Communications 17 (1): 5929.

Giotti, Bruno, Komal Dolasia, William Zhao, et al. 2024. “Single-Cell View of Tumor Microenvironment Gradients in Pleural Mesothelioma.” Cancer Discovery 14 (11): 2262–78.

Gorelick, Alexander N., Minsoo Kim, Walid K. Chatila, et al. 2021. “Respiratory Complex and Tissue Lineage Drive Recurrent Mutations in Tumour mtDNA.” Nature Metabolism 3 (4): 558–70.

Hart, Samuel F. M., Nicolas Alcala, Alison F. Feder, and Kelley Harris. 2026. “A Signature-Agnostic Test for Differences between Tumor Mutation Spectra Reveals Carcinogen and Ancestry Effects.” Communications Biology 9 (1): 462.

Hmeljak, Julija, Francisco Sanchez-Vega, Katherine A. Hoadley, et al. 2018. “Integrative Molecular Characterization of Malignant Pleural Mesothelioma.” Cancer Discovery 8 (12): 1548–65.

Ho, Mitchell, Raffit Hassan, Jingli Zhang, et al. 2005. “Humoral Immune Response to Mesothelin in Mesothelioma and Ovarian Cancer Patients.” Clinical Cancer Research: An Official Journal of the American Association for Cancer Research 11 (10): 3814– 20.

Huang, Junjie, Sze Chai Chan, Wing Sze Pang, et al. 2023. “Global Incidence, Risk Factors, and Temporal Trends of Mesothelioma: A Population-Based Study.” Journal of Thoracic Oncology: Official Publication of the International Association for the Study of Lung Cancer 18 (6): 792–802.

Imieliński, M., and X. Yao. 2024. “Mskilab/gGnome: gGnome: Reference Based Assembly Graph for Analyzing Rearranged Genomes.” Preprint, May 8. https://rdrr.io/github/mskilab/gGnome/#vignettes.

Imielinski, Marcin, Guangwu Guo, and Matthew Meyerson. 2017. “Insertions and Deletions Target Lineage-Defining Genes in Human Cancers.” Cell 168 (3): 460–472.e14.

Kim, Hoon, Nam-Phuong Nguyen, Kristen Turner, et al. 2020. “Extrachromosomal DNA Is Associated with Oncogene Amplification and Poor Outcome across Multiple Cancers.” Nature Genetics 52 (9): 891–97.

Koboldt, Daniel C., Qunyuan Zhang, David E. Larson, et al. 2012. “VarScan 2: Somatic Mutation and Copy Number Alteration Discovery in Cancer by Exome Sequencing.” Genome Research 22 (3): 568–76.

Korbel, Jan O., and Peter J. Campbell. 2013. “Criteria for Inference of Chromothripsis in Cancer Genomes.” Cell 152 (6): 1226–36.

Landrum, Melissa J., Jennifer M. Lee, Mark Benson, et al. 2018. “ClinVar: Improving Access to Variant Interpretations and Supporting Evidence.” Nucleic Acids Research 46 (D1): D1062–67.

Leonard, Jeanne, Marie Duhamel, and Michel Salzet. 2026. “Humoral Immunity in Cancer.” Signal Transduction and Targeted Therapy 11 (1): 335.

Liu, Gang, Paul Cheresh, and David W. Kamp. 2013. “Molecular Basis of Asbestos-Induced Lung Disease.” Annual Review of Pathology 8 (1): 161–87.

Love, Michael I., Wolfgang Huber, and Simon Anders. 2014. “Moderated Estimation of Fold Change and Dispersion for RNA-Seq Data with DESeq2.” Genome Biology 15 (12): 550.

Lu, Xiaofan, Jialin Meng, Yujie Zhou, Liyun Jiang, and Fangrong Yan. 2021. “MOVICS: An R Package for Multi-Omics Integration and Visualization in Cancer Subtyping.” Bioinformatics (Oxford, England) 36 (22–23): 5539–41.

Maclachlan, Kylee H., Even H. Rustad, Andriy Derkach, et al. 2021. “Copy Number Signatures Predict Chromothripsis and Clinical Outcomes in Newly Diagnosed Multiple Myeloma.” Nature Communications 12 (1): 5172.

Mangiante, Lise, Nicolas Alcala, Alexandra Sexton-Oates, et al. 2023. “Multiomic Analysis of Malignant Pleural Mesothelioma Identifies Molecular Axes and Specialized Tumor Profiles Driving Intertumor Heterogeneity.” Nature Genetics 55 (4): 607–18.

Mansfield, Aaron S., Tobias Peikert, James B. Smadbeck, et al. 2019. “Neoantigenic Potential of Complex Chromosomal Rearrangements in Mesothelioma.” Journal of Thoracic Oncology: Official Publication of the International Association for the Study of Lung Cancer 14 (2): 276–87.

Marshall, Kelly, Susan Jackson, Jennifer Jones, et al. 2020. “Homozygous Deletion of CDKN2A in Malignant Mesothelioma: Diagnostic Utility, Patient Characteristics and Survival in a UK Mesothelioma Centre.” Lung Cancer (Amsterdam, Netherlands) 150 (December): 195–200.

Martínez-Jiménez, Francisco, Peter Priestley, Charles Shale, Jonathan Baber, Erik Rozemuller, and Edwin Cuppen. 2023. “Genetic Immune Escape Landscape in Primary and Metastatic Cancer.” Nature Genetics 55 (5): 820–31.

Massey, Sheersh, Mohammad Aasif Khan, Safia Obaidur Rab, et al. 2023. “Evaluating the Role of MEN1 Gene Expression and Its Clinical Significance in Breast Cancer Patients.” PloS One 18 (7): e0288482.

McKenna, Aaron, Matthew Hanna, Eric Banks, et al. 2010. “The Genome Analysis Toolkit: A MapReduce Framework for Analyzing next-Generation DNA Sequencing Data.” Genome Research (United States) 20 (9): 1297–303.

Meiller, Clément, François Montagne, Theo Z. Hirsch, et al. 2021. “Multi-Site Tumor Sampling Highlights Molecular Intra-Tumor Heterogeneity in Malignant Pleural Mesothelioma.” Genome Medicine 13 (1): 113.

Nair, Nishanth Ulhas, Qun Jiang, Jun Stephen Wei, et al. 2023. “Genomic and Transcriptomic Analyses Identify a Prognostic Gene Signature and Predict Response to Therapy in Pleural and Peritoneal Mesothelioma.” Cell Reports. Medicine 4 (2): 100938.

Ng, Pauline C., and Steven Henikoff. 2003. “SIFT: Predicting Amino Acid Changes That Affect Protein Function.” Nucleic Acids Research 31 (13): 3812–14.

Nguyen, Luan, John W. M. Martens, Arne Van Hoeck, and Edwin Cuppen. 2020. “Pan-Cancer Landscape of Homologous Recombination Deficiency.” Nature Communications 11 (1): 1–12.

Nik-Zainal, Serena, Ludmil B. Alexandrov, David C. Wedge, et al. 2012. “Mutational Processes Molding the Genomes of 21 Breast Cancers.” Cell 149 (5): 979–93.

Qi, Meifang, Jiali Pang, Irene Mitsiades, Andrew A. Lane, and Esther Rheinbay. 2023. “Loss of Chromosome Y in Primary Tumors.” Cell 186 (14): S0092-8674(23)00646–3.

Reyniès, Aurélien de, Marie-Claude Jaurand, Annie Renier, et al. 2014. “Molecular Classification of Malignant Pleural Mesothelioma: Identification of a Poor Prognosis Subgroup Linked to the Epithelial-to-Mesenchymal Transition.” Clinical Cancer Research: An Official Journal of the American Association for Cancer Research 20 (5): 1323–34.

Rigutto, Angelica, Nicolás G. Núñez, Jenny C. Kienzler, et al. 2026. “Multiplexed Single-Cell and Spatial Profiling Reveal B Cells and Tertiary Lymphoid Structures as Prognostic Indicators in Pleural Mesothelioma.” British Journal of Cancer 135 (2): 214–22.

Seth, Sohan, and Manuel J. A. Eugster. 2016. “Probabilistic Archetypal Analysis.” Machine Learning 102 (1): 85–113.

Severson, David T., Assunta De Rienzo, and Raphael Bueno. 2020. “Mesothelioma in the Age of ‘Omics’: Before and after The Cancer Genome Atlas.” Journal of Thoracic and Cardiovascular Surgery 160 (4): 1078–1083.e2.

Severson, David T., Samuel Freyaldenhoven, Benjamin Wadowski, et al. 2026. “Multiomic, Histologic, and scRNA-Seq Profiling of Pleural Mesothelioma Reveals Negative Prognosis Associated with a Novel Uncommitted Molecular Phenotype.” Journal of Thoracic Oncology: Official Publication of the International Association for the Study of Lung Cancer 21 (4): 103529.

Shukla, Arti, Michael Jung, Maria Stern, et al. 2003. “Asbestos Induces Mitochondrial DNA Damage and Dysfunction Linked to the Development of Apoptosis.” American Journal of Physiology. Lung Cellular and Molecular Physiology 285 (5): L1018–25.

Sondka, Zbyslaw, Sally Bamford, Charlotte G. Cole, Sari A. Ward, Ian Dunham, and Simon A. Forbes. 2018. “The COSMIC Cancer Gene Census: Describing Genetic Dysfunction across All Human Cancers.” Nature Reviews. Cancer 18 (11): 696–705.

Soneson, Charlotte, Michael I. Love, and Mark D. Robinson. 2015. “Differential Analyses for RNA-Seq: Transcript-Level Estimates Improve Gene-Level Inferences.” F1000Research 4 (December): 1521.

Steele, Christopher D., Ammal Abbasi, S. M. Ashiqul Islam, et al. 2022. “Signatures of Copy Number Alterations in Human Cancer.” Nature 606 (7916): 984–91.

Tyagi, Ethika, Bin Liu, Chelsea Li, Tong Liu, Jared Rutter, and Douglas Grossman. 2017. “Loss of p16INK4A Stimulates Aberrant Mitochondrial Biogenesis through a CDK4/Rb-Independent Pathway.” Oncotarget 8 (34): 55848–62.

Van Loo, Peter, Silje H. Nordgard, Ole Christian Lingjærde, et al. 2010. “Allele-Specific Copy Number Analysis of Tumors.” Proceedings of the National Academy of Sciences of the United States of America 107 (39): 16910–15.

Wang, Hao, Tianwang Guan, Rong Hu, et al. 2025. “Targeting KAT7 Inhibits the Progression of Colorectal Cancer.” Theranostics 15 (4): 1478–95.

Wang, Lin, Magdalena Cybula, Maria Rostworowska, et al. 2022. “Upregulation of Succinate Dehydrogenase (SDHA) Contributes to Enhanced Bioenergetics of Ovarian Cancer Cells and Higher Sensitivity to Anti-Metabolic Agent Shikonin.” Cancers 14 (20): 5097.

Wang, Yan, Sufang Chen, Wei Tian, et al. 2019. “High-Expression HBO1 Predicts Poor Prognosis in Gastric Cancer.” American Journal of Clinical Pathology 152 (4): 517–26.

Watts, Stephen. 2024. Github.com/nf-core/oncoanalyser. Zenodo. 10.5281/ZENODO.16414049.

Woerner, August E., Jennifer Churchill Cihlar, Utpal Smart, and Bruce Budowle. 2020. “Numt Identification and Removal with RtN!” Bioinformatics (Oxford, England) 36 (20): 5115– 16.

Yu, Jixiang, Nanjun Chen, Zetian Zheng, Ming Gao, Ning Liang, and Ka-Chun Wong. 2023. “Chromothripsis Detection with Multiple Myeloma Patients Based on Deep Graph Learning.” Bioinformatics (Oxford, England) 39 (7): btad422.

Yuan, Yuan, Young Seok Ju, Youngwook Kim, et al. 2020. “Comprehensive Molecular Characterization of Mitochondrial Genomes in Human Cancers.” Nature Genetics 52 (3): 342–52.

